# Sense of self impacts spatial navigation and hexadirectional coding in human entorhinal cortex

**DOI:** 10.1101/2020.09.13.295246

**Authors:** Hyuk-June Moon, Baptiste Gauthier, Hyeong-Dong Park, Nathan Faivre, Olaf Blanke

## Abstract

Grid cells in entorhinal cortex (EC) encode an individual’s location in space and rely on environmental cues and self-motion cues derived from the individual’s body. Body-derived signals are also primary signals for the sense of self and self-identification with a body. However, it is currently unknown whether self-identification impacts grid cells and spatial navigation. Integrating the online manipulation of bodily signals, to modulate self-identification, into a spatial navigation task and with an fMRI measure to detect grid cell-like representation (GCLR) in humans, we report improved spatial navigation performance and decreased GCLR in EC when participants navigated with enhanced self-identification. These changes were further associated with increased activity in posterior parietal and retrosplenial cortex. Modulating self-identification by controlling online body-derived signals during spatial navigation, these data link the sense of self to spatial navigation and to entorhinal grid cell-like activity within a distributed cortical spatial navigation system.

## Introduction

The discovery of grid cells in rodent entorhinal cortex (EC) has shed new light on the neural mechanisms of spatial representation^1,2^. Grid cells are place-modulated neurons believed to represent the location of an individual and are defined by characteristic spatial firing field maps corresponding to hexagonal grid patterns that tile a given environment^3,4^. Entorhinal grid cell activity is modulated by sensory cues from the environment as well as by motion-related cues from the individual (i.e. self-motion cues)^1,2,5,6^. While the field maps of grid cells have been shown to depend on distal landmarks and field boundaries^1,7^, the periodic field maps of grid cells are maintained in darkness and across different environments and landmark changes^1,8^, suggesting movement-related input from the individual to grid cells. Subsequent studies demonstrated the primary importance of such self-motion cues from the individual’s body for generating and maintaining grid representations^9–11^. Overall, these findings support the proposal that grid cells keep track of an individual’s location in space by relying on both selfmotion cues and environmental sensory information^5,12^.

Self-motion cues are body-derived cues based on sensory and motor signals from the individual’s body during spatial navigation and include proprioceptive, tactile, vestibular, and motor signals^2,5,13^. Under normal conditions, the self is bound to the location of the physical body and experienced at the place occupied by the body. This association between self and body is a central feature of self-consciousness, captured by the concept of bodily self-consciousness (BSC)^14–16^. However, recent research using virtual reality (VR) has shown that illusory BSC states such as illusory self-identification can be induced for an avatar or virtual body employing the online manipulation of sensory and motor signals from the individual’s body^15,17^. Thus, during the full-body illusion^18,19^, participants view an avatar as seen from behind and projected in front of them, while an experimenter concurrently applies strokes to their back. Viewing one’s avatar and feeling one’s own back while being stroked synchronously, at these two different locations in space, leads to changes in self-identification with the avatar. Recent BSC studies, using the full-body illusion, have demonstrated that such bodily stimulations in VR, also impact tactile perception^20^, pain perception^21^, and temperature regulation^22^ . They also alter egocentric spatial processes including size perception^23–25^ and spatial semantic distance^26^, linking BSC with spatial processing. For example, Canzoneri et al. (2016) showed that when participants self-identified with a virtual body placed at a spatial location that differs from that of their physical body, the reference frame the brain used to compute abstract concepts (spatial semantical distance), which normally refers to the physical body, was shifted towards the position of the avatar. However, whereas the key importance of body-derived sensorimotor signals in grid cells is well documented, it is currently unknown whether and how online sensorimotor bodily stimulations in VR, that have been shown to modulate BSC, impact entorhinal grid cell activity. Here, we sought to investigate whether sensorimotor signals that alter BSC signals modulate grid cell activity in EC during spatial navigation in VR. While human grid cells have only rarely been described using single-unit recordings in epilepsy patients^27,28^, a method based on functional magnetic resonance imaging (fMRI) detecting a specific pattern in parametric BOLD (Blood-Oxygen-Level-Dependent) signal changes, the so-called *grid cell-like representation* (GCLR), has been proposed to reflect the activity of human grid cell populations^29–34^. GCLR is assumed to capture BOLD activity from populations of conjunctive grid cells in human EC^35^, characterized by headingdirection-dependent BOLD signal modulation with six-fold rotational symmetry following hexagonal grids of grid cells. Thus, the magnitude of hexadirectional BOLD signal modulation, GCLR, has been suggested as a proxy grid cell activity in humans^5,29,30^.

To investigate whether BSC impacts grid cell activity in EC, we designed a sensorimotor VR task and manipulated BSC while our participants performed a classical spatial navigation task as used in previous fMRI research^29,30^ that allowed us to assess spatial navigation performance and calculate GCLR. We hypothesized that experimentally induced changes in BSC would modulate GCLR, predicting that GCLR would be altered when enhanced BSC processing for the avatar is modulated by online sensorimotor stimulation. We, thus, assessed the impact of BSC on grid cells by systematically manipulating our participants’ self-identification with an avatar shown during navigation. Corroborated by behavioral data, our fMRI results show decreased GCLR in EC when participants navigated during an enhanced BSC state (i.e. navigating with a self-identified avatar). Increased BSC was further associated with increased BOLD activity in posterior parietal cortex, a core BSC region important for the integration of sensory and motor signals, and in retrosplenial cortex (RSC), a region mediating between ego- and allocentric spatial representation, collectively linking the sense of self to entorhinal grid cell activity and a distributed spatial navigation system.

## Results

### Avatar-related changes in BSC enhance spatial navigation performance

We adopted a spatial navigation task from previous fMRI studies^29,30^ to assess spatial navigation performance and BSC, and to calculate GCLR (Fig. 1a; see methods). Each session started with an encoding phase, in which participants had to memorize the locations of three objects. Following encoding, for each trial, a cue indicated a specific target object that participants had to recall and reach by navigating to it in the arena. At the end of each retrieval trial, the distance between the recalled location and its correct location (i.e. ‘distance error’) was determined. Navigation trace length and navigation time were also recorded in order to quantify spatial navigation performance. To assess the influence of BSC (self-identification) on grid cell-like activity as reflected in GCLR, we designed two experimental conditions and induced different levels of self-identification with the avatar by providing different online sensorimotor stimulation during the task. In the Body condition, supine participants saw, from their first-person viewpoint, a supine virtual avatar, which was spatially congruent with their own body position. As shown in Fig. 1a, we also showed the virtual right hand of the avatar (and a virtual joystick) that carried out the same movements as the participant’s right hand on the physical joystick in the scanner. Such spatial congruency between the participant’s body and the avatar’s body (or alignment of bodily reference frames; see methods for further detail) has been investigated in previous BSC research and shown to induce higher levels of self-identification with the avatar^36,37^. By contrast, the No-body condition did neither contain an avatar nor the right-hand movements and served as a control condition, for which we expected no or less self-identification as compared to the Body condition. Importantly, the No-body condition is identical to most previous human GCLR studies^29,30^.

**Fig. 1.**
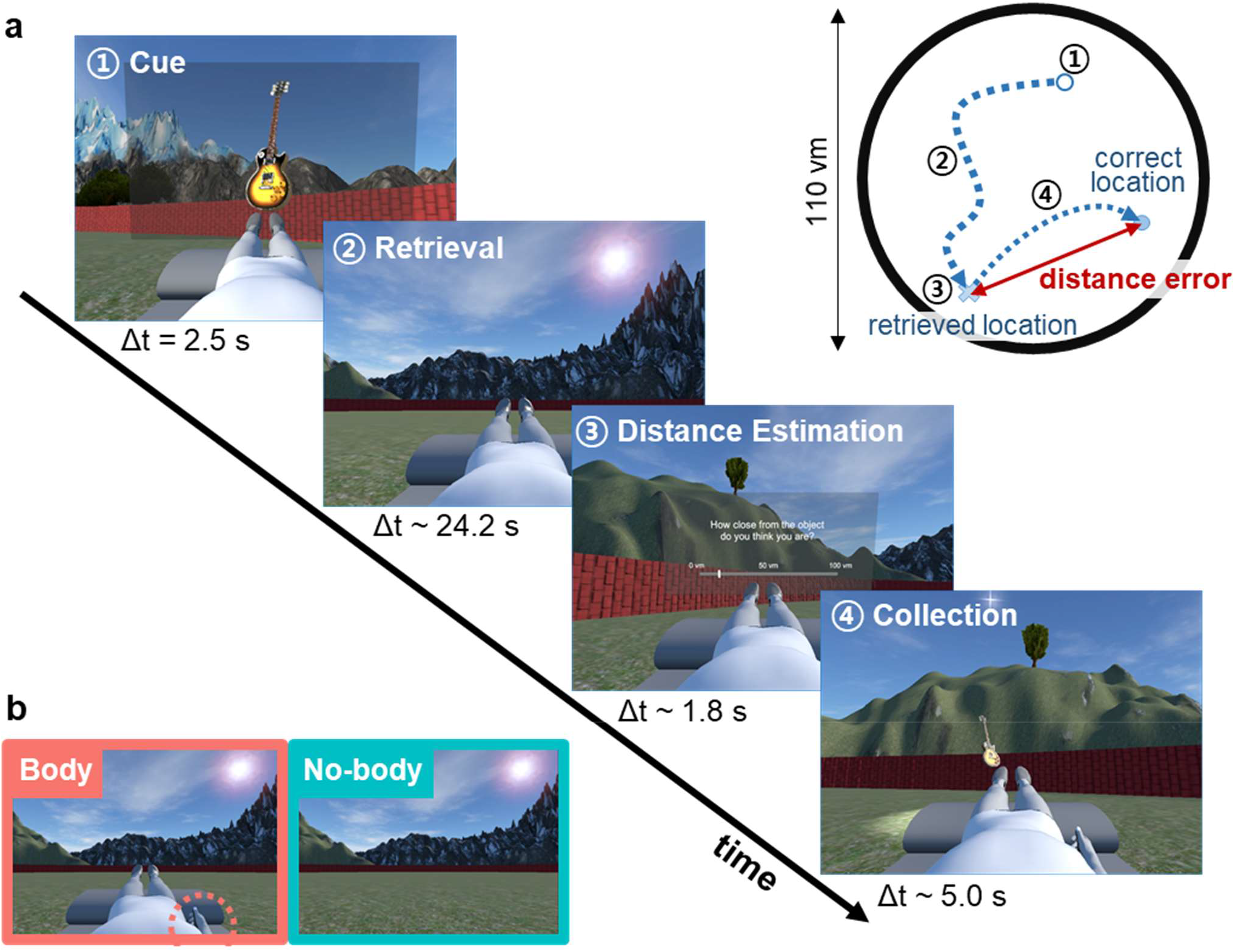
Spatial navigation task and Experimental BSC Conditions. **a**, The spatial navigation task consisted of six sessions with two experimental conditions. Each session started with an encoding phase, in which participants had to memorize the locations of three objects. Following encoding, participants performed 14 trials with the following steps: (1) Cue: a target object was provided (2.5 sec); (2) Retrieval: they had to recall and reach the original object location (self-paced, average 24.2 sec); (3) Distance estimation: they estimated the distance error they committed (self-paced, average 1.8 sec); (4) Collection: a target object appeared at its original location and participants were asked to navigate to it (self-paced, average 5.0 sec). **b**, In the Body condition, a body-shaped avatar (congruent with the posture and hand motion of the participant in the scanner) was seen by participants as part of the virtual scene during the entire procedure. In the No-body condition, the same scenes were displayed, but without the avatar (as is usually done during spatial navigation studies). Δt: mean duration, vm: virtual meter.

We assessed BSC by asking participants to rate their self-identification with the avatar (Q1: Self-Identification), to rate experienced threat (in response to a virtual knife that was seen as approaching the part of the arena where the virtual avatar was located) (Q2: Threat; see methods and supplementary Figure 1-c), and also assessed two control items (Q3, Q4; see methods). As predicted, ratings to Q1 and Q2 were higher in the Body vs. No-body condition (paired two-sided Wilcoxon signed-rank test, Q1: Z = 3.02, r = 0.60, p = 2.53e-03; Q2: Z = 3.80, r = 0.76, p = 1.42e-04, n = 25; Fig. 2a), indicating that our manipulation was effective in modulating BSC. Our participants did not give different ratings in the control questions, making it unlikely that the observed effects reflect the general response biases of the participants (i.e. no difference across conditions in control questions; Q3: Z = 0.57, r = 0.11, p = 0.569 Q4: Z = 1.05, r = 0.21, p = 0.293; see Methods; Supplementary Figure 1a). Post-experiment debriefing confirmed these results (see Methods; Supplementary Figure 1b).

**Fig. 2.**
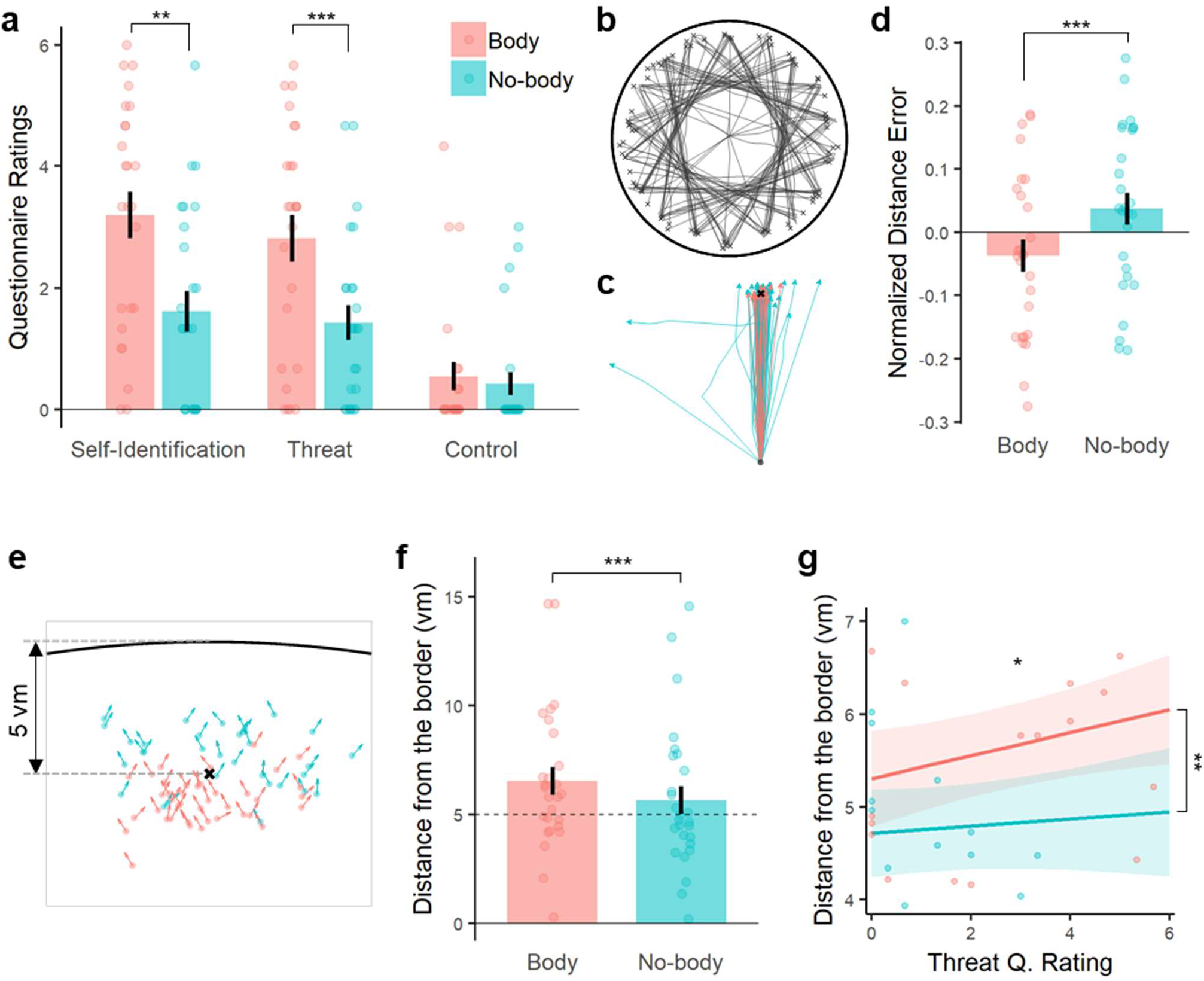
Enhanced self-identification was improved spatial navigation performance in the Body condition. **a**, Ratings of the questionnaire confirmed the effect of the experimental modulation on BSC. Self-identification (Q1) and experienced threat (Q2) were rated significantly higher in the Body vs. the No-body condition. **b**, Exemplary traces from a participant during the spatial navigation task. **c**, Overlay of the navigation traces per condition during the retrieval phase of the same participant (traces were rotated and shifted according to the starting and the target location, in order to better visualize the difference in distance errors and navigation efficiency). **d**, Participants showed better **spatial memory precision**, indexed by lower distance errors from the correct retrieval targets. In the graph, distance errors were z-scored within-participants for visualization, while the statistical analysis was performed with the raw values through a dedicated mixed model. Each dot represents the mean of individual participant per condition. **e**, The arrows display the trial-by-trial reached locations and heading directions of an exemplary participant. Locations are plotted relative to the correct target point (‘x’), and the arena’s border (black bold line). **f**, Participants stopped farther away from the arena’s border compared to the No-body condition during which no virtual avatar was presented. The condition-wise difference in the distance from the border was consistent across participants (22 out of 27). For the figure, participant-wise mean distance errors were visualized, while the statistical analysis was performed with the trial-wise values through a dedicated mixed model. **g**, Mixed-effect model slopes relating Threat (Q2) to the distance from the border in the two conditions, while taking into account the condition-wise difference. *: 0.01 <= p < 0.05, ** : 0.001 <= p < 0.01, *** : p < 0.001. The error bar indicates standard error.

To assess the influence of the BSC modulation on spatial navigation and memory, we compared spatial navigation/memory performance during both conditions. All participants were able to navigate in the virtual environment and complete the task while being scanned in the MRI scanner (Fig. 2b). Interestingly, participants showed better spatial memory precision as indexed by lower distance errors in the Body vs. No-body condition (mixed-effects regression; df = 1, F = 11.18, p = 8.25e-04, n = 27; Fig. 2d). Navigation efficiency was also improved, as participants carried out shorter paths in the Body condition vs. the No-body condition (df = 1, F = 8.46, p = 3.81e-03; Supplementary Figure 2a), while average navigation time did not differ across conditions (df = 1, F = 0.21, p = 0.648; Supplementary Figure 2b). These results demonstrate that the Body condition induces higher self-identification with the virtual avatar (as indexed by Q1 and Q2), as well as higher spatial navigation/memory performance in the virtual environment.

Of note, participants stopped navigating significantly farther from the arena’s border in the Body condition (i.e. the condition where they see a self-identified avatar in front of them) compared to the No-body condition (df = 1, F = 52.29, p = 4.80e-13, n = 27; Fig. 2e,f). This navigational difference was consistently observed in 22 out of 27 participants. Our finding is compatible with spatial changes, referred to as a drift in self-location toward a self-identified avatar, reported by previous research on BSC using different behavioral measures^18–20,38^. Thus, when self-identifying with the avatar seen in front of them, our participants stopped before reaching the intended destination, in turn, farther from the border (Fig. 2e). This link between the drift in spatial navigation and BSC was further confirmed by the significant relation between the drift and BSC ratings (i.e. Threat; Q2): the more participants felt threatened by the virtual knife directed to where the avatar was, the farther they stopped away from the arena’s border before reaching the target (1.00 ± 0.27 vm; predicted by mixed-effects regression; df = 1, F = 11.23, p = 0.0263, n = 25; Fig. 2g). We reiterate that spatial memory precision was higher in the Body condition, despite the fact that, on average, participants stopped farther from the optimal target point (5 vm) in the Body condition (6.38 ± 0.70 vm away from the border; Fig. 2f-red) compared to the No-body condition (5.38 ± 0.67 vm; Fig. 2f-blue). Thus, although it could be argued that the drift may worsen the distance error in the Body condition (participants stopped too early), the opposite was the case for overall spatial navigation performance. Angular errors were not affected by this drift effect and were also significantly lower in the Body condition (mixed-effects regression; df = 1, F = 5.397, p = 0.020, n = 27; Supplementary Figure 2c). To summarize, these behavioral results show that the Body condition was characterized by higher self-identification with the avatar and drift in self-location influencing navigational behavior, and, importantly, by an improvement in spatial navigation performance.

### Grid cell-like representation decreases when spatial navigation is performed with self-identified avatar

We next assessed whether these changes in BSC and spatial navigation were associated with changes in grid cell activity as reflected by GCLR in EC. As a first step, we confirmed the recruitment of GCLR in our task regardless of the experimental condition, applying previously established methods^29,31,34^. A putative grid-orientation (*φ*) of each session was estimated with the other five sessions of fMRI images matched with heading direction (*θ*) information. Based on the calculated grid-orientation, GCLR of each session was determined by the magnitude of the six-fold symmetric fluctuation as a function of the heading direction (see methods). This analysis revealed a significant hexadirectional BOLD signal modulation, GCLR, in EC when our participants navigated in the virtual environment (sinusoidal regressor: Z = 2.23, r = 0.45, p = 0.0128, Fig. 3b; aligned vs. misaligned contrast: Z = 2.00, r = 0.40, p = 0.0226, n = 25, Fig. 3c), replicating earlier data^29^.

**Fig. 3.**
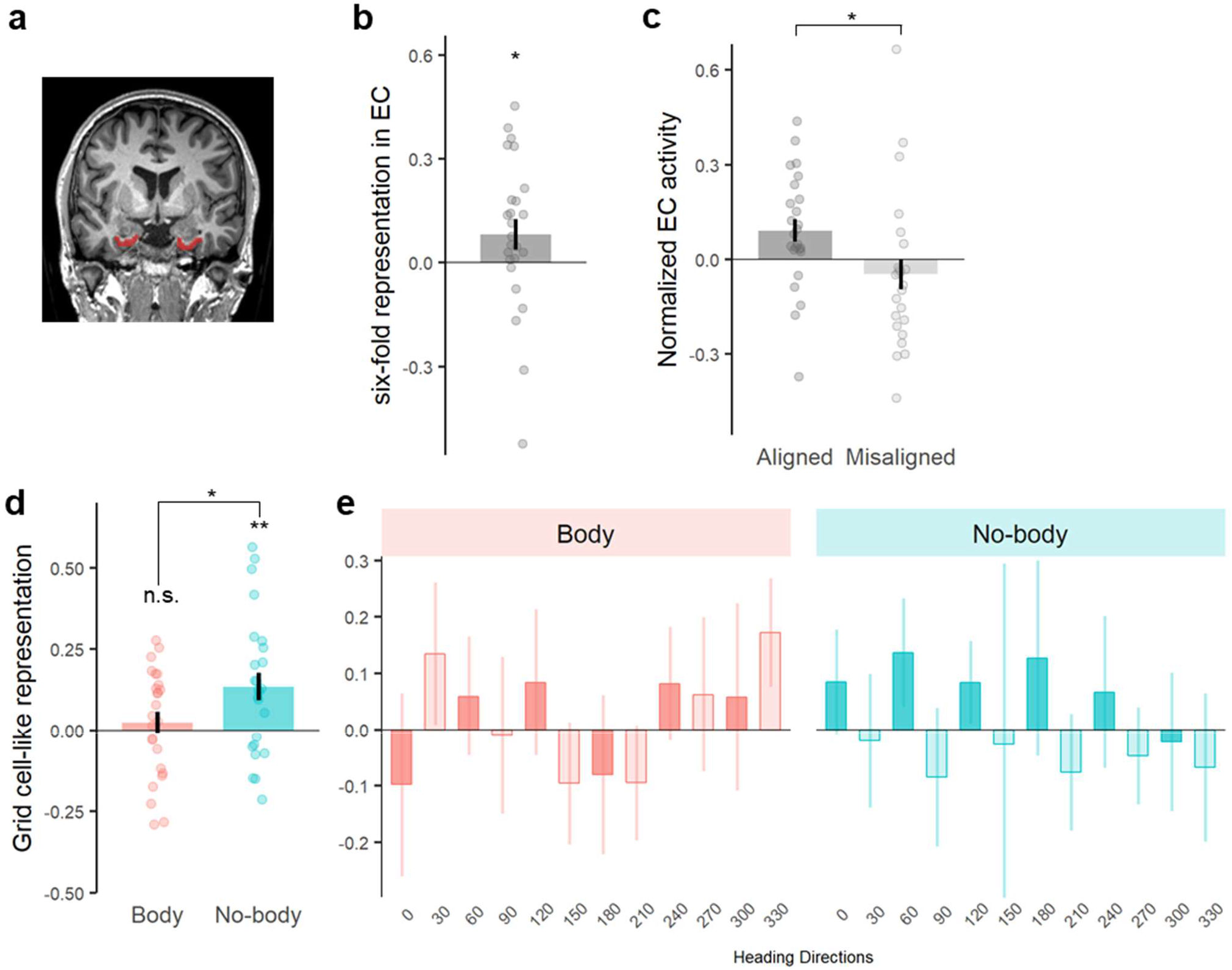
Decrease of entorhinal grid cell-like representation in Body condition. **a**, EC ROI of an exemplary participant **b**, Significant six-fold symmetric grid cell-like representation (GCLR) in human entorhinal cortex (EC), calculated for the entire recording session regardless of the experimental condition. **c**, EC activity during aligned navigation was significantly higher than during the misaligned navigation. d, Conditionwise GCLRs were significantly higher in the No-body (standard spatial navigation condition) than the Body condition. Notably, condition-wise grid cell-like representations in the Body condition were not significantly greater than zero, implying that the difference between conditions can be attributed to a reduced GCLR in the Body condition. **e**, Normalized EC activity profiles for every 30° heading direction showed that the typical hexadirectional modulation was hardly observable in the Body condition (left), while it was prominent in the No-body condition (right). n.s. : p >= 0.05, *: 0.01 <= p < 0.05, ** : p < 0.01. The error bar indicates standard error.

Next, we determined GCLR differences between the two conditions by calculating conditionwise GCLR using cross-validation, which was designed to be independent of the other sessions and, in turn, each condition(see Methods). These results confirmed that GCLR was pronounced in the No-body condition (the condition that is similar to conditions used by previous human spatial navigation and GCLR studies; Z = 2.83, r = 0.57, p = 2.30e-03, n = 25; Fig. 3d). GCLR was absent in the Body condition (Z = 0.69, r = 0.14, p = 0.245; Fig. 3d), which is the condition with enhanced BSC and spatial navigation performance. Within-subject comparisons between both conditions confirmed significantly lower GCLR in the Body vs. Nobody condition (Z = 2.26, r = 0.45, p = 0.0236).

To further investigate what might have led to these changes in GCLR across both conditions, we determined whether the aligned versus misaligned activity contrast of each voxel in the EC was indeed attenuated or whether the spatial and/or temporal stability of the putative grid orientation was merely deteriorated and veiled the existing hexadirectional modulation (see Methods). These analyses revealed that the average amplitude of voxel-wise GCLR was significantly lower in the Body condition and that the grid orientations in the Body condition were both spatially and temporally less stable (although these were not found to significantly differ between conditions; see Supplementary Results and Supplementary Figure 4). These additional data support our main fMRI result that – when performing spatial navigation with enhanced BSC – the GCLR is attenuated and hardly detectable, mainly based on decreased amplitudes of the voxel-level hexadirectional modulation in EC.

### RSC activity correlates with improved spatial navigation performance

In order to investigate the brain systems possibly accounting for the improved spatial navigation performance in the Body condition, we first assessed the correlation between GCLR and spatial memory precision. However, this was not found to be significant (df = 1, F = 0.022, p = 0.717, n = 25). Even though the grid cell system in EC is known to play a key role in spatial navigation^5,39^, previous human spatial navigation studies showed that other brain regions, such as retrosplenial cortex (RSC) and parahippocampal gyrus (PHC), are also prominently involved and often closely associated with spatial navigation performance^40–46^. To investigate this in other potentially involved brain regions, we applied whole-brain fMRI analysis (generalized linear model, GLM) and detected five clusters showing significant task-related activations (independently of the experimental conditions), which included the bilateral RSC, bilateral PHC, and right lingual gyrus (LiG) (Fig. 4a, Supplementary Figure 5a, Supplementary Table 2), consistently with the existing spatial navigation literature. Comparing activity in each of these five regions of interest (ROIs) between the Body vs. No-body condition during the task phases determining spatial memory precision (i.e. Cue and Retrieval, Fig. 1a), we observed significantly greater activity in right RSC (Fig. 4b; Z = 2.65, r = 0.53, p = 0.040, n = 25; Bonferroni-corrected). No significant differences were found in any of the other four regions (bilateral PHC, left RSC, right LiG; Supplementary Figure 5b). We further observed that higher right RSC activation was associated with better spatial memory precision (characterized by a smaller distance error; Fig. 2d) (df = 1, F = 12.11, p = 0.024, n = 24; Fig. 4c), further linking right RSC to improved spatial navigation performance in the Body condition. The results reveal the prominent implication of RSC in the present task and its contribution to improved spatial navigation performance in the Body condition.

**Fig. 4.**
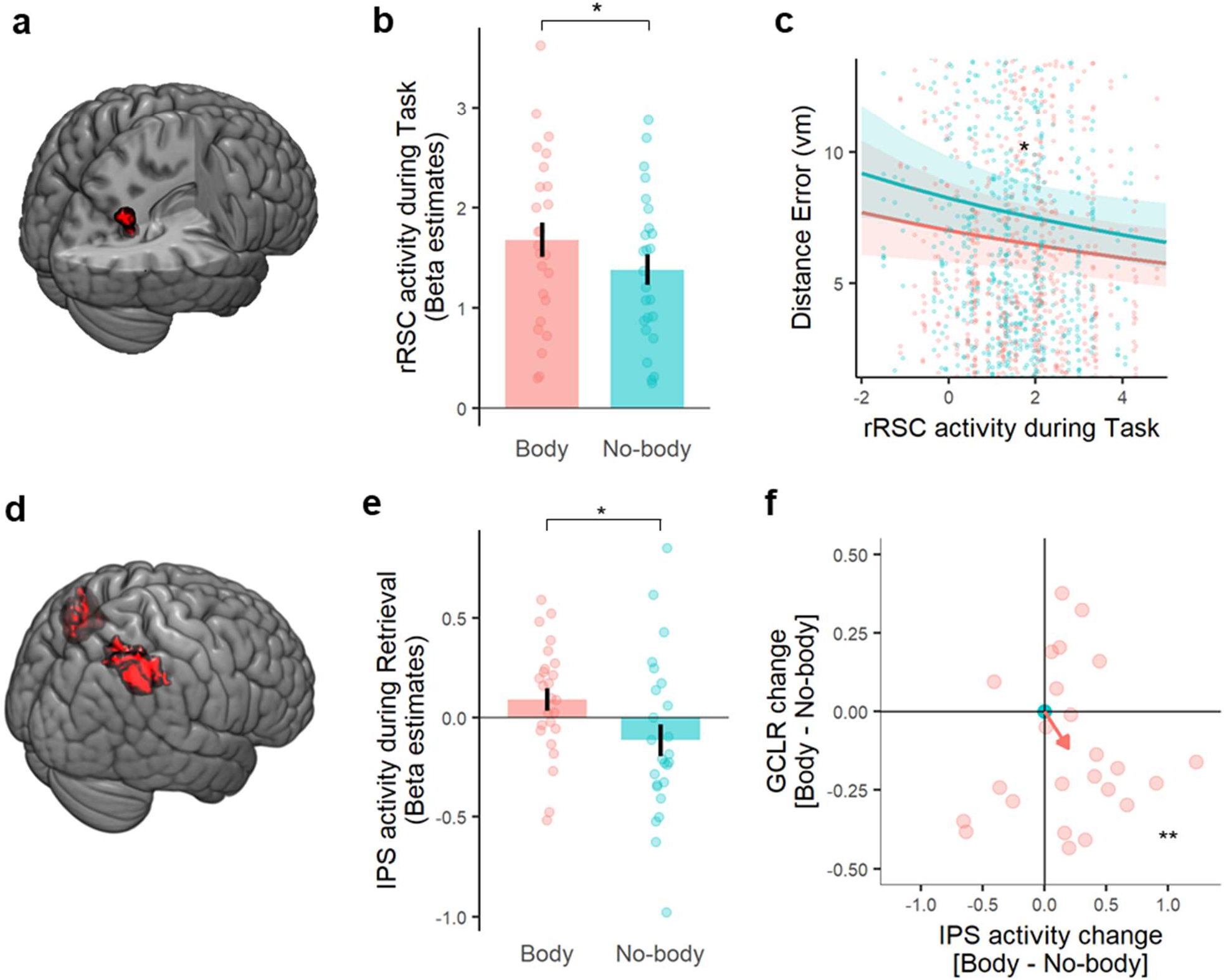
Retrosplenial cortex (RSC) and intraparietal sulcus (IPS) activity were increased in the Body condition. **a**, Functional localizer revealed that the right RSC was involved in the spatial navigation task. **b**, ROI analysis showed that right RSC was significantly more activated during the task (Bonferroni-corrected for five task-related clusters). **c**, The higher right RSC activity during the task phase before they reach the recalled location (i.e. Cue & Retrieval Phase) could predict better spatial memory precision. The depicted distance error range does not cover all data points as the full range plot (0~ 110 vm) cannot properly demonstrate the relationship between distance error and right RSC activity. **d**, Anatomical display of the a priori IPS ROI arguably activated during egocentric processing in link with BSC. **e**, IPS activity is significantly greater during navigation in the Body condition, where sensorimotor bodily signal integration takes place when participants are manipulating the joystick to navigate. This suggests that the experimental modulation of BSC boosted egocentric processes especially relevant to integrating sensorimotor bodily signals. **f**, Participant-wise IPS activity changes and GCLR changes in the Body condition with respect to the No-body condition. The plot demonstrates that performing the task with a self-identified avatar reduced GCLR while strengthening the IPS activity (multinomial test: p = 5.4e-03, post-hoc binomial test: p = 3.1e-03, n = 24). The red arrow indicates mean changes across participants. **\*** : 0.01 < p < 0.05, ** : p < 0.01. The error bar indicates standard error.

### Posterior parietal cortex activity is enhanced during spatial navigation in the Body condition

In an additional control analysis, we assessed whether a core region of BSC, the intraparietal sulcus region (IPS) in posterior parietal cortex^14,15,47–49^, was differently involved in the two spatial navigation conditions. Importantly, IPS is a core region not only for the integration of multisensory bodily signals and BSC, but also for egocentric spatial processing in spatial navigation^44,50,51^. We expected that IPS activity (Fig. 4d; see Methods) would be enhanced in the BSC-enhanced Body condition and that this would be especially the case during navigation because only during navigation did participants receive different online sensorimotor signals (i.e. while participants are navigating by manipulating the joystick). In accordance with our expectation, we found significantly greater IPS activation during navigation (i.e. Retrieval phase) in the Body vs. No-body condition (Fig. 4e; Z = 2.44, r = 0.49, p = 0.0147, n = 25), but not during the Cue phase (Z = 0.85, r = 0.17, p = 0.396). Participant-wise changes in IPS activity, as well as GCLR in EC, demonstrated that IPS activity was increased in the Body vs. No-body condition, while GCLR was decreased in the Body (Fig. 4f). These associated changes, induced by the BSC modulation (i.e. self-identification with the avatar), thereby link both structures: IPS and EC. This was found again only during navigation (multinomial test: p = 5.4e-03, post-hoc binomial test with H0 probability 0.25: p = 3.1e-03, n = 24), but not during the non-navigation cue phase (multinomial test; p = 0.09). These IPS and GCLR results associate GCLR attenuation, during spatial navigation with a self-identified avatar, with increased IPS activation in the same condition.

## Discussion

Here we show that signals that are of relevance for self-consciousness contribute to grid cell-like activity by associating an enhanced state of BSC during spatial navigation in a virtual environment with improved performance in a spatial navigation task and decreased GCLR in EC, which has been proposed to reflect the activity of grid cell populations^29–34^. These data link entorhinal grid cell-like activity to the sense of self, as based on online body-derived sensorimotor signals applied during spatial navigation. Enhanced BSC was further associated with increased activation in posterior parietal cortex and retrosplenial cortex, showing that experimentally-controlled changes in the sense of self impact cortical spatial navigation system beyond EC.

### Entorhinal GCLR and BSC

Entorhinal grid cell activity in animals and humans has been consistently shown to depend on self-motion-related cues, originating from the individual’s body, as well as sensory cues from the environment^1,2,5,6^. Prior human studies have also observed GCLR linked to various cognitive functions related to spatial representation^29,32–34,52,53^. However, it is not known how GCLR in humans depends on the sense of self and on BSC in particular. In the present study, we observed the typical hexadirectional modulation in EC during the condition that was similar to previous spatial navigation studies (i.e. the No-body condition)^29–31^, while the GCLR in EC was attenuated in the Body condition, linking GCLR reduction to enhanced BSC. Decreases in GCLR could result not only from the reduced amplitude of heading direction-dependent BOLD signal variations in EC (regardless of a putative grid orientation), but may also stem from the instability of the grid orientations either in time or across voxels in the ROI (even if head direction-dependent BOLD modulations are strong)^29–31^. In-depth analysis of our results (Supplementary Figure S4 and Supplementary Results) shows that the voxel-wise amplitudes of the hexadirectional modulation were lower in the Body vs. No-body condition, while the stabilities of grid orientations did not differ significantly. Accordingly, we argue that the reduced GCLR observed in the Body condition was likely due to a decrease in the amplitude of voxel-level hexadirectional BOLD modulations, compatible with an overall reduction of the activity of the grid cell system in the EC, rather than a discord of active, but not co-aligned, grid cells. Further results from the present study suggest that the decreased GCLR is not attributable to navigation-related factors (such as speed, central navigational preference, differences in target locations, and training) nor to head-motion artifacts, changes in attention, or visual occlusion by the avatar (for a detailed discussion see Supplementary discussion).

The decreased GCLR observed in the Body condition could simply be a consequence of a decoupling between the spatial navigation behavior and the grid cell system, thus reflecting the flexibility of its involvement in the spatial representation rather than its impairment ^30,31^. Indeed, a recent human study showed that the theta oscillation in the medial temporal lobe encoding the boundary information does not present when the navigation was not relevant to the target search task^54^, suggesting the dynamic involvement of the sub-systems in the medial temporal lobe . More relevant to our study, it has been also reported that spatial navigation based on non-allocentric strategy does not recruit the hippocampal-entorhinal regions, while activation of the area is observed during navigation based on allocentric strategies^55^. Therefore, we suggest that the reduction of GCLR in the Body condition could be related to comparable mechanisms: the enhanced self-identification with the avatar could boost egocentric processing and indirectly decreasing the reliance on allocentric strategies. Interestingly, previous findings have shown that the reference point during egocentric spatial processing is not necessarily the participant’s physical body per se, but at a location, where the participant experiences to be located in space (under most conditions, of course, the body’s position, but see ^26,56^). This is also consistent with our RSC and IPS data which are discussed further below.

We report that sensorimotor congruency between participants’ physical body and the seen corresponding avatar during spatial navigation improved spatial navigation performance: participants committed smaller distance errors while navigating in shorter paths when seeing the avatar associated with synchronous sensorimotor stimulation (Fig. 2c,d). Previous work has revealed that modulations of BSC not only alter body- and self-related processes^20–22^, but also affect spatial representation and episodic memory^24,26,57,58^. The present data also extend previous BSC research to the field of spatial navigation and grid cells, successfully modulating BSC by enhancing self-identification with the avatar during spatial navigation. The *explicit* BSC changes, measured by questionnaires, were further linked with *implicit* behavioral BSC changes characterized by a drift in self-location. Participants stopped farther away from the arena’s border when navigating with a self-identified avatar (as compared to the No-body condition), revealing a BSC change that altered where our participants located themselves with respect to external landmarks. These novel self-location findings extend previous BSC data using gait responses^18,20^, mental imagery of spatial distance^19,59^, or imagined spatial navigation^38^ to the field of spatial navigation (for review, see ^38,59–61^) and confirm our hypothesis that the sense of self affects processes of spatial navigation. The present self-location and spatial navigation data suggest that the reference point during spatial navigation in the Body condition was processed with respect to the avatar in the VR space, linking enhanced BSC and sensorimotor processing centered on the avatar with improved spatial navigation performance.

The suggested decoupling between GCLR and spatial navigation in the Body condition could also explain the observed improvement in spatial navigation performance despite GCLR reduction that seems to contradict the known role of grid cells in spatial navigation^5,39,62^. Congruent with our suggestion, we did not detect a correlation between GCLR and spatial memory performance, as was also not the case in many previous human grid cell studies using similar spatial navigation paradigms^29,31,33,63^. While some GCLR studies reported a positive correlation between GCLR and spatial memory performance^29,64^, different parameters and neuroimaging modalities were used for these correlations. Importantly, not all studies used the same spatial navigation paradigm, which may greatly affect the involvement of grid cell system (e.g. pre-training, ego- vs. allocentric strategy). Hence, it is difficult to generalize from the results of these different studies to our results. Finally, it has been shown that place cells also encode the locations of others^65,66^ and this may also apply to grid cells as tested here for GCLR with an avatar that we showed in a virtual environment. Human GCLR has also been reported during conceptual spatial mapping^32^, and thus manipulations not related to the position of the navigating agent per se. Accordingly, one cannot directly assume that stronger self-identification with respect to an avatar in a virtual environment leads to enhanced GCLR.

It is also possible that the absence of GCLR in the Body condition originated from the deformation of grid cells’ grids, as the GCLR is calculated on the premise of the six-fold symmetric grid firing fields map. For instance, in line with rodent data^67^, physical restrictions of navigation possibilities in the environment (e.g. by physical spatial constraints) have been shown to disrupt the hexadirectional GCLR modulation in humans^68^. In the presence of such restrictions, firing fields of grid cells do not tile in typical hexagonal grids, reflecting a change of navigational processing corresponding to the dimensions of the navigation corridor. Besides, distortion of grid cells’ spatial representation has also been linked to the integration of incongruent visual and self-motion cues in rodents ^11^. Here the authors showed that grid cell firing field maps can be elongated or shrunk by the relationship of proprioceptive versus visual cues and that the distorted grid patterns are no longer six-fold symmetric. In our study, the two conditions were performed in the same virtual arena, thus there should be no influence of physical restrictions. However, changes in self-identification were induced by fully-controlled sensorimotor stimulation that linked the physical body with the virtual avatar. It is thus also possible that such stimulations may have interfered with the integration across different bodily reference frames during navigation (i.e. visual environment-avatar versus participant’s body in the scanner), thereby leading to the observed GCLR decrease. Although we cannot exclude this as an alternative explanation, we do not think that it is very likely because the conflict between the reference frames always exists in virtual navigation paradigms, yet it was rather reduced in our controlled experiment. Moreover, the self-motion cues provided in the Body condition (i.e. hand motions) were not directly relevant to spatial navigation and, thus, grid cells’ firing fields. However, we agree that these important issues need further investigation in humans, for instance through single neuron recordings.

### RSC and PPC in spatial navigation and BSC

EC is not the sole brain region that determines spatial navigation performance. For example, Kunz et al. (2015) reported that compensatory mechanisms in hippocampus account for GCLR reduction despite maintained spatial navigation performance. Spatial navigation is based on activity within a large distributed network, involving IPS/PPC, RSC, and several other regions^42–44,46,69–71^, also supported by whole-brain analysis in the present study revealing activations in bilateral retrosplenial cortex (RSC), bilateral parahippocampal gyrus and right lingual gyrus^40–43^. From these regions, only RSC activity was enhanced in the Body condition and we further associated higher RSC activation with improved spatial navigation performance (i.e. smaller distance error). Previous work consistently linked human RSC activation to spatial navigation^43,45,72–76^ and RSC has been proposed to be important for orienting to landmarks^63^, a central feature of our experimental design as distal landmarks only provided orientation cues. Moreover, clinical research consistently linked human RSC damage (especially of right RSC) to orientation impairments in spatial navigation^45,77,78^. Hence, the correlation we observed between RSC activation and spatial memory precision (i.e. distance error) extends previous findings about the role of RSC in spatial navigation and adds the important novel finding that spatial navigation-related activity in RSC, depends on the level of BSC. RSC has been regarded as a mediator between self-centered (egocentric) and environmental (allocentric) processes in PPC and medial EC, respectively^72,73^. RSC integrates body-derived self-motion cues while mapping one’s location in the environment^79,80^. Moreover, RSC has also prominently been associated with several “self”-related cognitive processes beyond spatial navigation, such as “self”-orientation in time^81^, across social dimensions^82^, integration of “self”-referential stimuli^83^, autobiographical memory^84,85^, and BSC^15,47,48^.

The present IPS data further extend the BSC-related changes we observed in RSC and EC. IPS, and more generally PPC, is regarded as a core region for egocentric spatial representation in humans^44,50,86,87^. Supporting this human work, rodent studies reported that neurons in PPC encode self-centered cues independent of the external environment (e.g. selfmotion and acceleration) during navigation^88,89^. In addition, many human BSC studies reported IPS activation when key components of BSC (e.g. self-location and self-identification) were modulated by multisensory stimuli, and IPS is considered a key BSC region^14,15,47–49^. Reporting increased IPS activity associated with solid behavioral evidence of drift in self-location during spatial navigation when our participants navigated with a self-identified avatar, these IPS and RSC data extend previous BSC neuroimaging findings to the field of spatial navigation.

In the Body condition, we observed increased IPS activation that has been linked with BSC and egocentric spatial processing. Many previous studies showed that body-referenced cues (e.g. vestibular, motor, and proprioceptive signals) are processed in PPC and provide crucial inputs to grid cells^10,11,27,90^. We argue that the present data link BSC-related processing as manipulated by online sensorimotor stimulation to egocentric processes in IPS and to allocentric grid cells in EC, suggesting that human grid cell-like activity in EC reflects ego-versus allocentric processing demands. Enhanced RSC activity in the Body condition and the mediating role of RSC between ego- and allocentric processes in spatial navigation, as well as between PPC and EC^72,73^, is compatible with this suggestion. Accordingly, we propose that the BSC changes, characterized by strengthened “self”-centered processing referenced to the avatar, associate enhanced egocentric, self-centered, processing in IPS, with altered ego- and allocentric processing in RSC, and with reduced allocentric spatial representation, GCLR in EC.

## Conclusion

By linking an attenuated GCLR in EC with an enhanced BSC state induced by online sensorimotor stimulation while participants performed a spatial navigation task, we demonstrate that experimentally-controlled changes in BSC modulate activity in the grid cell system. Both systems (grid cells, BSC) rely on body-derived signals as a primary input to encode an individual’s location in space. The present data show that the human grid cell system is influenced by sensorimotor BSC-signals that associate the participant’s physical body with a virtual body during navigation. Increased BSC was further associated with increased activation in IPS, a core BSC region important for the integration of sensory and motor signals, and RSC, a region mediating between ego- and allocentric spatial representation. Collectively, the present data link entorhinal grid cell activity with BSC and show that experimentally-controlled changes in the sense of self impact GCLR in EC and a cortical spatial navigation system beyond EC.

## Methods

### Participants

Twenty-seven healthy right-handed participants (13 males and 14 females; mean age 25.3±1.96) with normal or corrected-to-normal vision were recruited from the general population. The number of participants, 27, was chosen according to the minimum sample size calculated from Nau et al. (2018) to reproduce conventional grid cell-like representation. Participants were naive to the purpose of the study, gave informed consent in accordance with the institutional guidelines (IRB #: GE 15-273) and the Declaration of Helsinki (2013), and received monetary compensation (CHF20/hour). Two participants who entered random answers to the questionnaire excluded from the questionnaire analysis (i.e. both pressed the response button repeatedly at incorrect moments during the experiment; further confirmed by post-experiment debriefing). A participant whose structural image was examined as abnormal by a medical investigator was excluded from the fMRI analysis. Another participant was excluded from the grid cell-like representation analyses due to severe image distortions and signal drop in EC (~2.3 % of voxels in the EC were above the global average of mean EPI). A session with head drift greater than 3 mm was also excluded from fMRI analyses.

### MRI-compatible Virtual Reality (VR) spatial navigation task

Stereoscopic visual stimuli were provided via MRI-compatible goggles (VisualSystem, Neuro Nordic Lab: 800×600 resolution, 40Hz refresh rate). An MRI-compatible joystick (Tethyx, Current Designs) was used to perform the task inside the MRI scanner. The task program including the virtual arena and virtual object was implemented with Unity Engine (Unity Technologies, https://unity3d.com).

The task arena did not contain any landmark inside, and distal landmarks providing orientation cues were placed outside of its boundary. The task procedures were adopted from previous human spatial navigation studies (Fig. 1a)^29,30,33^. Each session of the task started with an encoding phase, in which participants had to sequentially and repeatedly memorize the locations of three objects (at least three times per object), while freely navigating inside the circular arena using a joystick. In each trial, following encoding, participants were asked to recall and return to where a cued object was. (1) During the Cue phase, one target object among the three encoded objects was represented as floating in the virtual scene for 2 s. (2) After the cue disappeared, they had to navigate to the target location, where they recalled the target object was placed before, by manipulating the joystick. After reaching the recalled position, they pressed a button on the joystick to confirm their response (i.e. Retrieval phase). (3) Sequentially, participants were asked to report the distance error they estimated to have committed (the distance between the reported and correct object’s location; Fig. 1a-top-right) by indicating on a continuous scale ranging from 0 to 110 vm (i.e. Distance-estimation phase). (4) Following the distance estimation, the object appeared at its correct position, and participants had to navigate and collect the object (i.e., Collection phase) before starting the next trial. The Collection phase was to provide a participant with an additional encoding cue (i.e. feedback), but also to ensure that the spatial traces spanned various directions designed to allow the analysis of grid cell-like representation (GCLR) (Fig. 2b; Horner et al., 2016). At the end of each session, participants were presented with a virtual knife directed towards them, in order to measure how threatened they felt as a subjective measure of BSC change (Fig. 2a-Q2; see Questionnaire section).

In total, the experiment consisted of six sessions divided into three blocks, aimed at comparing the two conditions (Fig. 1b) in terms of both brain activity and spatial navigation performances. The order of the conditions was pseudo-randomized within each block and counterbalanced between participants (N-B/B-N/N-B or B-N/N-B/B-N). As the task was self-paced, the duration of each session varied depending on the participant’s performance (mean round duration: 9.0 ± 0.70 min), but didn’t differ between the conditions (t-test: p = 0.92).

### BSC modulation with the virtual body

The Body condition with a neutral body-shaped avatar was designed to experimentally modulate BSC, more specifically, to enhance self-identification with the virtual avatar in the VR environment compared to the baseline: the No-body condition. The avatar was designed to be gender-neutral and grey-skinned without hair. The virtual body was in the same posture as a participant – supine – and its right hand was shown as congruent with respect to the participant’s hand movements controlling the joystick. These settings of avatar were chosen to achieve sensorimotor congruency between the participant’s body and the avatar, which has been reported to lead to self-identification with the avatar^36,37,91,92^.

### Questionnaire

At the end of each session, participants were asked to answer four questions using a Likert scale ranging from −3 (strongly disagree) to 3 (strongly agree). The questions were randomly ordered across sessions and participants answered with the joystick. Q1 (“I felt as if what I saw in the middle of the scene was my body”) was intended to measure self-identification with the avatar. Q2 (“I felt as if the threat (knife) was toward me”) was also designed to measure the degree of threat towards the participant. Q3 (“I felt dizzy”) sought to measure cybersickness (Supplementary Figure 1a). Q4 (“I felt as if I had 3 bodies”) served as a general control question. A short debriefing was carried out after participants had completed the experiment.

### Prescreening and training in the Mock scanner

The participants were trained to perform the spatial navigation task in a mock scanner. The training consisted of one session of the No-body condition and lasted around ten minutes, keeping them naive to the experimental condition. To avoid potential carryover effects, we used different virtual objects and environment than those used in the main experiment. This training also allowed us to exclude participants experiencing a severe cybersickness caused by navigation in the VR environment^93^.

### MRI data acquisition

MRI data were acquired at the Human Neuroscience Platform of the Campus Biotech (Geneva, Switzerland), with a 3T MRI scanner (SIEMENS, MAGNETOM Prisma) equipped with a 64-channel head-and-neck coil. The task-related functional images covering the entire brain were acquired with a T2*-weighted Echo Planar Imaging (EPI) sequence with the following parameters: TR = 1000 ms, TE = 32 ms, Slice thickness = 2mm (no gap), In-plane resolution = 2 mm x 2 mm, Number of slices = 66, Multiband factor = 6, FoV = 225 mm, Flip angle = 50°, slice acquisition order = interleaved. The structural image per participant was recorded with a T1-weighted MPRAGE sequence with the following parameters: TR = 2300 ms, TE = 2.25 ms, TI = 900ms, Slice thickness = 1mm, In-plane resolution = 1 mm x 1 mm, Number of slices = 208, FoV = 256 mm, Flip angle = 8°. In addition to that, B0 field map (magnitude and phase information, respectively) was acquired to correct EPI distortion by inhomogeneous magnetic fields (especially, near the medial temporal lobe).

### fMRI data preprocessing

MRI data were preprocessed with SPM12 (http://www.fil.ion.ucl.ac.uk/spm). Functional images were slice-time corrected, realigned, unwarped using B0 field map and co-registered with the individual T1-weighted structural image. For the conventional generalized linear model(GLM) analysis (e.g. region-of-interest(ROI) analysis of task-related regions and IPS), the images were normalized to Montreal Neurological Institute (MNI) space, to allow a second-level GLM analysis designed to localize commonly activated brain regions across participants. Following previous studies, other analyses regarding grid cell-like hexadirectional modulation with the EC as a main ROI were conducted in the native space without normalization to avoid additional signal distortion^30,31^. All preprocessed functional images were smoothed with a 5mm full-width-half-maximum Gaussian smoothing kernel as the final preprocessing step.

### Definition of the EC ROI in participants’ native space

Participant-wise EC ROIs for the analysis of grid cell-like representations(GCLR) were defined using Freesurfer (v6.0.0, http://surfer.nmr.mgh.harvard.edu) as described in previous studies^30,94^. Briefly, a cortical parcellation was automatically conducted by the software with the individual T1 structural images based on the Desikan-Killiany Atlas^95^. The bilateral EC labels generated from the parcellation were taken as an individual ROI and were examined manually by overlapping them on the corresponding structural image. Subsequently, the ROIs in the ‘freesurfer conformed space’ were transformed into volume ROIs in the participant’s native space and, again, ‘coregistered and resliced’ to the mean EPI images.

### Analysis of grid-cell like representation (GCLR)

The Grid Code Analysis Toolbox (GridCAT v1.03, https://www.nitrc.org/projects/gridcat) under MATLAB 2018b (The Mathworks) was used to analyze GCLR^96^, following a seminal method which was proposed by Doeller et al. (2010). The analysis was comprised of two steps with mutually exclusive datasets. As a first step, with one of the partitioned datasets, a first GLM (GLM 1) was used to calculate *β*_1_ and *β*_2_, using two parametric modulation regressors: *cos*(6*θ_t_*) and *sin*(6*θ_t_*) respectively, where *θ_t_* is heading direction during the navigation in time(t). Then, using the betas, voxel-wise amplitude(*A*) and grid orientations(*φ*) were respectively estimated by 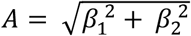 and *φ* = *tan*^-1^(*β*_2_/*β*_1_) /6.

In the second part of the analysis, a putative grid orientation was calculated by the weighted average of the voxel-wise grid orientations(*φ*) in the ROI (i.e. EC) with the voxel-wise amplitude(*A*) of each voxel as its weight. Subsequently, based on the putative grid-orientation(*φ*) and moving direction information(*θ_t_*), a second GLM (GLM2) estimates an amplitude of GCLR in the EC, selectively (1) by contrasting regressors for navigation toward grid-aligned (*φ* + 0,60,120, …, 300) vs. misaligned (*φ* + 30,90,150, …, 330) direction or (2) by applying a six-fold symmetric sinusoidal parametric modulation regressor: *cos*(6(*θ_t_* – *φ*)). We calculated GCLRs with both methods respectively, to confirm that our result does not depend on either of the methods. In order to optimally utilize all available data and improve the signal-to-noise ratio, a cross-validation method with multiple partitions was adopted^34^, instead of dividing the data into two equal halves. An amplitude of six-fold representation for a given session was calculated based on the grid orientation estimated with the other five sessions and the process was repeated for every session. The session-wise results were summed together to estimate the overall grid cell-like representation of the participant.

### Calculation of condition-wise GCLR

The classical GCLR in the previous step was calculated with all six sessions without taking into account the experimental condition of each session, which implies that each estimate does not purely represent a magnitude of the hexadirectional modulation in the corresponding experimental condition. Besides, as two different experimental conditions were timely intermingled, grid orientations across sessions even within the same condition could be unstable, which critically affects the GCLR estimation^30,31^. Hence, we calculated session-wise BOLD contrasts between aligned vs. misaligned movement independently from the other sessions, using data from the single session only. Leave-one-out cross-validation with ten partitions was performed for each session to optimally utilize the dataset and maximize the signal to noise ratio (See the Methods section above)^34^. Notably, this method was dedicated to the comparison of GCLR between the Body and No-body condition, rather than the demonstration of hexadirectional modulation in contrast to the controls (e.g. four/five/seven-fold symmetry) which was already fulfilled. Calculated session-wise results were averaged by condition to get the condition-wise GCLR, which are robust to potential temporal instability of grid orientations across sessions.

### Temporal and spatial stabilities of grid orientations

Spatial stability was defined as the homogeneity of voxel-wise grid orientations within EC. To assess the spatial stability of a session, Rayleigh’s test for non-uniformity of circular data was calculated with the voxel-wise grid orientations within EC. Rayleigh’s z-value was taken as an index of the spatial stability of the session^30,31^. Temporal stability was defined as the stability of grid orientations over time. For each session, it was computed by the circular standard deviation of ten grid orientations estimated during each of the ten cross-validations of GCLR described above. Of note, the ten grid orientations were calculated with ten different data portions from different time points. Therefore temporally stable grid orientations should remain similar across folds, resulting in a small standard deviation.

### Generalized linear model (GLM) analysis to detect task-related brain regions

Whole-brain GLM analysis using normalized function images was performed to detect brain regions, possibly accounting for changes in spatial navigation performance. Beta values during the task phase were extracted using GLM analysis with the tailored regressors using SPM12 (Supplementary Table 1). First, with the parametric modulation regressor, we searched for brain regions where its activation during the ‘Retrieval’ phase was correlated with the ‘distance error’. However, this analysis did not reveal any significant clusters after the voxel-level family-wise error correction (FWE). Second, we assessed contrasts (Body > Nobody) during the task phases before participants finished the retrieval procedure (i.e. Cue + Retrieval), as such responses could be responsible for spatial navigation performance of the trial. However, again, no cluster (extent threshold > 20 voxels) survived after voxel-level FWE correction (p < 0.05).

### ROI analysis of task-related BOLD activity by using the functional localizer

Next, we performed ROI analysis to investigate brain regions involved in the spatial navigation performance. Functional ROIs relevant to the spatial navigation task were defined by the functional contrast (Task: Cue + Retrieval + Feedback + Collection > implicit baseline; i.e. orthogonal to the conditions of interest: the Body and No-body) in 2^nd^ Level GLM using a voxel-level threshold family-wise error-corrected for multiple comparisons of p < 0.05 and using an extent threshold of 20 voxels (Fig. 4a; Supplementary Figure 5; Supplementary Table 2). In order to search for task-related activity accounting for improved spatial navigation, mean beta values within the ROIs during the task phases before reaching the recalled location (i.e. Cue + Retrieval) were compared between the two experimental conditions.

### ROI analysis of intraparietal sulcus region (IPS)

We anatomically defined the ROI for the bilateral IPS based on the normalized functional images (in MNI space) by using SPM Anatomy Toolbox based on the Jülich probabilistic cytoarchitectonic maps^97^. Betas during the Retrieval phase in IPS were compared between the Body and No-body condition.

### Statistical analysis

Statistical assessments for behavioral and fMRI data were performed with R (v3.5.3 for Windows, https://www.r-project.org/) and RStudio (v1.2.1335, http://www.rstudio.com). Outlying data points outside of the standard deviation range, −3 to 3, were excluded from the statistical analysis. For the behavioral parameters having a value per trial (distance errors, navigation trace length and time, distance from the border), mixed-effects regressions (lme4_1.1-18-1, a package of R), which includes condition as a fixed effect and random intercepts for individual participants, were used to assess statistical significance. Random slopes were included as far as there were no estimation failures. For the other parameters with no single-trial estimates (e.g. questionnaire ratings, spatial/temporal stability of the grid orientation, task-related BOLD activity), non-parametric two-sided Wilcoxon signed-rank test was used. The conventional six-fold patterns of GCLR were assessed with a one-sided Wilcoxon signed-rank test, as expected from previous work^29,34^. However, the condition-wise comparison of the GCLRs was performed with a two-sided test. Assessments of correlations were conducted using mixed-effect regression models so that the condition-induced effects within participants are properly taken into account by the random effect of a participant, in addition to the across-participants effect. In order to assume the best-fit distribution and apply proper parameters for each mixed-effects regression used, data distribution of each dependent variable was assessed using fitdistrplus(v1.0-11, a package of R).

## Acknowledgments

This work was supported by the Korean Government Scholarship Program for study overseas and the Bertarelli Foundation to HM. OB is supported by the Swiss National Science Foundation (No. 320030_188798) and by the Bertarelli Foundation. Additional support by the Fondation Campus Biotech Geneva (FCBG) - a foundation of the Swiss Federal Institute of Technology Lausanne (EPFL), the University of Geneva (UniGe), and the Hôpitaux Universitaires de Genève (HUG), the Institute of Translational Molecular Imaging (ITMI).

## Author Contributions

Conceptualization, H.M., B.G, H.P., and O.B.; Methodology, H.M., B.G., H.P., and N.F.; Software, H.M.; Formal Analysis, H.M., and N.F.; Investigation, H.M.; Writing – Original Draft, H.M.; Writing – Review & Editing, B.G., H.P., N.F., and O.B.; Resources, H.M.; Funding Acquisition, O.B.; Supervision, O.B.

## Competing Interests

The authors declare no competing interests.

## Data and code availability

The data that support the findings of this study and the analysis code are available from the corresponding author upon reasonable request.

## Notes

### Competing Interest Statement

The authors have declared no competing interest.

### Summary of Updates

In this version of the manuscript, the discussion has been revised to reduce possible confusion. And, figures have been merged to reduce the number of the figures.

## References

1 Hafting, T., Fyhn, M., Molden, S., Moser, M. B. & Moser, E. I. Microstructure of a spatial map in the entorhinal cortex. Nature 436, 801–806, doi:10.1038/nature03721 (2005).

2 Moser, E. I., Kropff, E. & Moser, M. B. Place cells, grid cells, and the brain’s spatial representation system. Annu Rev Neurosci 31, 69–89, doi:10.1146/annurev.neuro.31.061307.090723 (2008).

3 Stemmler, M., Mathis, A. & Herz, A. V. Connecting multiple spatial scales to decode the population activity of grid cells. Sci Adv 1, e1500816, doi:10.1126/science.1500816 (2015).

4 Ismakov, R., Barak, O., Jeffery, K. & Derdikman, D. Grid Cells Encode Local Positional Information. Curr Biol 27, 2337–2343 e2333, doi:10.1016/j.cub.2017.06.034 (2017).

5 Rowland, D. C., Roudi, Y., Moser, M. B. & Moser, E. I. Ten Years of Grid Cells. Annu Rev Neurosci 39, 19–40, doi:10.1146/annurev-neuro-070815-013824 (2016).

6 Barry, C. & Burgess, N. Neural mechanisms of self-location. Curr Biol 24, R330–339, doi:10.1016/j.cub.2014.02.049 (2014).

7 Barry, C., Hayman, R., Burgess, N. & Jeffery, K. J. Experience-dependent rescaling of entorhinal grids. Nat Neurosci 10, 682–684, doi:10.1038/nn1905 (2007).

8 Fyhn, M., Hafting, T., Treves, A., Moser, M. B. & Moser, E. I. Hippocampal remapping and grid realignment in entorhinal cortex. Nature 446, 190–194, doi:10.1038/nature05601 (2007).

9 Jacob, P. Y., Poucet, B., Liberge, M., Save, E. & Sargolini, F. Vestibular control of entorhinal cortex activity in spatial navigation. Front Integr Neurosci 8, 38, doi:10.3389/fnint.2014.00038 (2014).

10 Campbell, M. G. et al. Principles governing the integration of landmark and self-motion cues in entorhinal cortical codes for navigation. Nat Neurosci 21, 1096–1106, doi:10.1038/s41593-018-0189-y (2018).

11 Chen, G., Lu, Y., King, J. A., Cacucci, F. & Burgess, N. Differential influences of environment and self-motion on place and grid cell firing. Nat Commun 10, 630, doi:10.1038/s41467-019-08550-1 (2019).

12 McNaughton, B. L., Battaglia, F. P., Jensen, O., Moser, E. I. & Moser, M. B. Path integration and the neural basis of the ‘cognitive map’. Nat Rev Neurosci 7, 663–678, doi:10.1038/nrn1932 (2006).

13 Britten, K. H. Mechanisms of self-motion perception. Annu Rev Neurosci 31, 389–410, doi:10.1146/annurev.neuro.29.051605.112953 (2008).

14 Park, H. D. & Blanke, O. Coupling Inner and Outer Body for Self-Consciousness. Trends Cogn Sci 23, 377–388, doi:10.1016/j.tics.2019.02.002 (2019).

15 Blanke, O., Slater, M. & Serino, A. Behavioral, Neural, and Computational Principles of Bodily Self-Consciousness. Neuron 88, 145–166, doi:10.1016/j.neuron.2015.09.029 (2015).

16 Blanke, O. Multisensory brain mechanisms of bodily self-consciousness. Nat Rev Neurosci 13, 556–571, doi:10.1038/nrn3292 (2012).

17 Tsakiris, M. My body in the brain: a neurocognitive model of body-ownership. Neuropsychologia 48, 703–712, doi:10.1016/j.neuropsychologia.2009.09.034 (2010).

18 Lenggenhager, B., Tadi, T., Metzinger, T. & Blanke, O. Video ergo sum: manipulating bodily self-consciousness. Science 317, 1096–1099, doi:10.1126/science.1143439 (2007).

19 Ionta, S. et al. Multisensory mechanisms in temporo-parietal cortex support self-location and first-person perspective. Neuron 70, 363–374, doi:10.1016/j.neuron.2011.03.009 (2011).

20 Aspell, J. E., Lenggenhager, B. & Blanke, O. Keeping in touch with one’s self: multisensory mechanisms of self-consciousness. PLoS One 4, e6488, doi:10.1371/journal.pone.0006488 (2009).

21 Hansel, A., Lenggenhager, B., von Kanel, R., Curatolo, M. & Blanke, O. Seeing and identifying with a virtual body decreases pain perception. Eur J Pain 15, 874–879, doi:10.1016/j.ejpain.2011.03.013 (2011).

22 Salomon, R., Lim, M., Pfeiffer, C., Gassert, R. & Blanke, O. Full body illusion is associated with widespread skin temperature reduction. Front Behav Neurosci 7, 65, doi:10.3389/fnbeh.2013.00065 (2013).

23 Banakou, D., Groten, R. & Slater, M. Illusory ownership of a virtual child body causes overestimation of object sizes and implicit attitude changes. Proc Natl Acad Sci U S A 110, 12846–12851, doi:10.1073/pnas.1306779110 (2013).

24 van der Hoort, B., Guterstam, A. & Ehrsson, H. H. Being Barbie: the size of one’s own body determines the perceived size of the world. PLoS One 6, e20195, doi:10.1371/journal.pone.0020195 (2011).

25 Pasqualini, I., Llobera, J. & Blanke, O. “Seeing” and “feeling” architecture: how bodily self-consciousness alters architectonic experience and affects the perception of interiors. Front Psychol 4, 354, doi:10.3389/fpsyg.2013.00354 (2013).

26 Canzoneri, E., di Pellegrino, G., Herbelin, B., Blanke, O. & Serino, A. Conceptual processing is referenced to the experienced location of the self, not to the location of the physical body. Cognition 154, 182–192, doi:10.1016/j.cognition.2016.05.016 (2016).

27 Jacobs, J. et al. Direct recordings of grid-like neuronal activity in human spatial navigation. Nat Neurosci 16, 1188–1190, doi:10.1038/nn.3466 (2013).

28 Nadasdy, Z. et al. Context-dependent spatially periodic activity in the human entorhinal cortex. Proc Natl Acad Sci U S A 114, E3516–E3525, doi:10.1073/pnas.1701352114 (2017).

29 Doeller, C. F., Barry, C. & Burgess, N. Evidence for grid cells in a human memory network. Nature 463, 657–661, doi:10.1038/nature08704 (2010).

30 Kunz, L. et al. Reduced grid-cell-like representations in adults at genetic risk for Alzheimer’s disease. Science 350, 430–433, doi:10.1126/science.aac8128 (2015).

31 Stangl, M. et al. Compromised Grid-Cell-like Representations in Old Age as a Key Mechanism to Explain Age-Related Navigational Deficits. Curr Biol 28, 1108–1115 e1106, doi:10.1016/j.cub.2018.02.038 (2018).

32 Constantinescu, A. O., O’Reilly, J. X. & Behrens, T. E. Organizing conceptual knowledge in humans with a gridlike code. Science 352, 1464–1468, doi:10.1126/science.aaf0941 (2016).

33 Horner, A. J., Bisby, J. A., Zotow, E., Bush, D. & Burgess, N. Grid-like Processing of Imagined Navigation. Curr Biol 26, 842–847, doi:10.1016/j.cub.2016.01.042 (2016).

34 Nau, M., Navarro Schroder, T., Bellmund, J. L. S. & Doeller, C. F. Hexadirectional coding of visual space in human entorhinal cortex. Nat Neurosci 21, 188–190, doi:10.1038/s41593-017-0050-8 (2018).

35 Kunz, L. et al. Mesoscopic Neural Representations in Spatial Navigation. Trends Cogn Sci 23, 615–630, doi:10.1016/j.tics.2019.04.011 (2019).

36 Walsh, L. D., Moseley, G. L., Taylor, J. L. & Gandevia, S. C. Proprioceptive signals contribute to the sense of body ownership. J Physiol 589, 3009–3021, doi:10.1113/jphysiol.2011.204941 (2011).

37 Kokkinara, E. & Slater, M. Measuring the effects through time of the influence of visuomotor and visuotactile synchronous stimulation on a virtual body ownership illusion. Perception 43, 43–58, doi:10.1068/p7545 (2014).

38 Nakul, E., Orlando-Dessaints, N., Lenggenhager, B. & Lopez, C. Measuring perceived self-location in virtual reality. Sci Rep 10, 6802, doi:10.1038/s41598-020-63643-y (2020).

39 Moser, M. B., Rowland, D. C. & Moser, E. I. Place cells, grid cells, and memory. Cold Spring Harb Perspect Biol 7, a021808, doi:10.1101/cshperspect.a021808 (2015).

40 Zhang, H. & Ekstrom, A. Human neural systems underlying rigid and flexible forms of allocentric spatial representation. Hum Brain Mapp 34, 1070–1087, doi:10.1002/hbm.21494 (2013).

41 Wolbers, T. & Buchel, C. Dissociable retrosplenial and hippocampal contributions to successful formation of survey representations. J Neurosci 25, 3333–3340, doi:10.1523/JNEUROSCI.4705-04.2005 (2005).

42 Boccia, M., Nemmi, F. & Guariglia, C. Neuropsychology of environmental navigation in humans: review and meta-analysis of FMRI studies in healthy participants. Neuropsychol Rev 24, 236–251, doi:10.1007/s11065-014-9247-8 (2014).

43 Ranganath, C. & Ritchey, M. Two cortical systems for memory-guided behaviour. Nat Rev Neurosci 13, 713–726, doi:10.1038/nrn3338 (2012).

44 Ekstrom, A. D., Arnold, A. E. & Iaria, G. A critical review of the allocentric spatial representation and its neural underpinnings: toward a network-based perspective. Front Hum Neurosci 8, 803, doi:10.3389/fnhum.2014.00803 (2014).

45 Epstein, R. A. Parahippocampal and retrosplenial contributions to human spatial navigation. Trends Cogn Sci 12, 388–396, doi:10.1016/j.tics.2008.07.004 (2008).

46 Maguire, E. A., Burgess, N. & O’Keefe, J. Human spatial navigation: cognitive maps, sexual dimorphism, and neural substrates. Curr Opin Neurobiol 9, 171–177, doi:10.1016/s0959-4388(99)80023-3 (1999).

47 Guterstam, A., Bjornsdotter, M., Gentile, G. & Ehrsson, H. H. Posterior cingulate cortex integrates the senses of self-location and body ownership. Curr Biol 25, 1416–1425, doi:10.1016/j.cub.2015.03.059 (2015).

48 Grivaz, P., Blanke, O. & Serino, A. Common and distinct brain regions processing multisensory bodily signals for peripersonal space and body ownership. Neuroimage 147, 602–618, doi:10.1016/j.neuroimage.2016.12.052 (2017).

49 Petkova, V. I. et al. From part-to whole-body ownership in the multisensory brain. Curr Biol 21, 1118–1122, doi:10.1016/j.cub.2011.05.022 (2011).

50 Byrne, P., Becker, S. & Burgess, N. Remembering the past and imagining the future: a neural model of spatial memory and imagery. Psychol Rev 114, 340–375, doi:10.1037/0033-295X.114.2.340 (2007).

51 Cabeza, R., Ciaramelli, E., Olson, I. R. & Moscovitch, M. The parietal cortex and episodic memory: an attentional account. Nat Rev Neurosci 9, 613–625, doi:10.1038/nrn2459 (2008).

52 Bellmund, J. L., Deuker, L., Navarro Schroder, T. & Doeller, C. F. Grid-cell representations in mental simulation. Elife 5, doi:10.7554/eLife.17089 (2016).

53 Julian, J. B., Keinath, A. T., Frazzetta, G. & Epstein, R. A. Human entorhinal cortex represents visual space using a boundary-anchored grid. Nat Neurosci 21, 191–194, doi:10.1038/s41593-017-0049-1 (2018).

54 Stangl, M. et al. Boundary-anchored neural mechanisms of location-encoding for self and others. Nature 589, 420–425, doi:10.1038/s41586-020-03073-y (2021).

55 Iaria, G., Petrides, M., Dagher, A., Pike, B. & Bohbot, V. D. Cognitive strategies dependent on the hippocampus and caudate nucleus in human navigation: variability and change with practice. J Neurosci 23, 5945–5952 (2003).

56 Noel, J. P., Pfeiffer, C., Blanke, O. & Serino, A. Peripersonal space as the space of the bodily self. Cognition 144, 49–57, doi:10.1016/j.cognition.2015.07.012 (2015).

57 Bergouignan, L., Nyberg, L. & Ehrsson, H. H. Out-of-body-induced hippocampal amnesia. Proc Natl Acad Sci U S A 111, 4421–4426, doi:10.1073/pnas.1318801111 (2014).

58 Brechet, L. et al. First-person view of one’s body in immersive virtual reality: Influence on episodic memory. PLoS One 14, e0197763, doi:10.1371/journal.pone.0197763 (2019).

59 Lenggenhager, B., Mouthon, M. & Blanke, O. Spatial aspects of bodily self-consciousness. Conscious Cogn 18, 110–117, doi:10.1016/j.concog.2008.11.003 (2009).

60 Dieguez, S. & Lopez, C. The bodily self: Insights from clinical and experimental research. Ann Phys Rehabil Med 60, 198–207, doi:10.1016/j.rehab.2016.04.007 (2017).

61 Heydrich, L. et al. Visual capture and the experience of having two bodies - Evidence from two different virtual reality techniques. Front Psychol 4, 946, doi:10.3389/fpsyg.2013.00946 (2013).

62 Buzsaki, G. & Moser, E. I. Memory, navigation and theta rhythm in the hippocampal-entorhinal system. Nat Neurosci 16, 130–138, doi:10.1038/nn.3304 (2013).

63 Bierbrauer, A. et al. Unmasking selective path integration deficits in Alzheimer’s disease risk carriers. Sci Adv 6, eaba1394, doi:10.1126/sciadv.aba1394 (2020).

64 Maidenbaum, S., Miller, J., Stein, J. M. & Jacobs, J. Grid-like hexadirectional modulation of human entorhinal theta oscillations. Proc Natl Acad Sci U S A 115, 10798–10803, doi:10.1073/pnas.1805007115 (2018).

65 Danjo, T., Toyoizumi, T. & Fujisawa, S. Spatial representations of self and other in the hippocampus. Science 359, 213–218, doi:10.1126/science.aao3898 (2018).

66 Omer, D. B., Maimon, S. R., Las, L. & Ulanovsky, N. Social place-cells in the bat hippocampus. Science 359, 218–224, doi:10.1126/science.aao3474 (2018).

67 Derdikman, D. et al. Fragmentation of grid cell maps in a multicompartment environment. Nat Neurosci 12, 1325–1332, doi:10.1038/nn.2396 (2009).

68 He, Q. & Brown, T. I. Environmental Barriers Disrupt Grid-like Representations in Humans during Navigation. Curr Biol 29, 2718–2722 e2713, doi:10.1016/j.cub.2019.06.072 (2019).

69 Barra, J., Laou, L., Poline, J. B., Lebihan, D. & Berthoz, A. Does an oblique/slanted perspective during virtual navigation engage both egocentric and allocentric brain strategies? PLoS One 7, e49537, doi:10.1371/journal.pone.0049537 (2012).

70 Igloi, K., Doeller, C. F., Berthoz, A., Rondi-Reig, L. & Burgess, N. Lateralized human hippocampal activity predicts navigation based on sequence or place memory. Proc Natl Acad Sci U S A 107, 14466–14471, doi:10.1073/pnas.1004243107 (2010).

71 Berthoz, A. & Viaud-Delmon, I. Multisensory integration in spatial orientation. Curr Opin Neurobiol 9, 708–712, doi:10.1016/s0959-4388(99)00041-0 (1999).

72 Mitchell, A. S., Czajkowski, R., Zhang, N., Jeffery, K. & Nelson, A. J. D. Retrosplenial cortex and its role in spatial cognition. Brain Neurosci Adv 2, 2398212818757098, doi:10.1177/2398212818757098 (2018).

73 Vann, S. D., Aggleton, J. P. & Maguire, E. A. What does the retrosplenial cortex do? Nat Rev Neurosci 10, 792–802, doi:10.1038/nrn2733 (2009).

74 Marchette, S. A., Vass, L. K., Ryan, J. & Epstein, R. A. Anchoring the neural compass: coding of local spatial reference frames in human medial parietal lobe. Nat Neurosci 17, 1598–1606, doi:10.1038/nn.3834 (2014).

75 Shine, J. P., Valdes-Herrera, J. P., Hegarty, M. & Wolbers, T. The Human Retrosplenial Cortex and Thalamus Code Head Direction in a Global Reference Frame. J Neurosci 36, 6371–6381, doi:10.1523/JNEUROSCI.1268-15.2016 (2016).

76 Committeri, G. et al. Reference frames for spatial cognition: different brain areas are involved in viewer-, object-, and landmark-centered judgments about object location. J Cogn Neurosci 16, 1517–1535, doi:10.1162/0898929042568550 (2004).

77 Maguire, E. A. The retrosplenial contribution to human navigation: a review of lesion and neuroimaging findings. Scand J Psychol 42, 225–238, doi:10.1111/1467-9450.00233 (2001).

78 Hashimoto, R., Tanaka, Y. & Nakano, I. Heading disorientation: a new test and a possible underlying mechanism. Eur Neurol 63, 87–93, doi:10.1159/000276398 (2010).

79 Elduayen, C. & Save, E. The retrosplenial cortex is necessary for path integration in the dark. Behav Brain Res 272, 303–307, doi:10.1016/j.bbr.2014.07.009 (2014).

80 Alexander, A. S. & Nitz, D. A. Retrosplenial cortex maps the conjunction of internal and external spaces. Nat Neurosci 18, 1143–1151, doi:10.1038/nn.4058 (2015).

81 Peer, M., Lyon, R. & Arzy, S. Orientation and disorientation: lessons from patients with epilepsy. Epilepsy Behav 41, 149–157, doi:10.1016/j.yebeh.2014.09.055 (2014).

82 Peer, M., Salomon, R., Goldberg, I., Blanke, O. & Arzy, S. Brain system for mental orientation in space, time, and person. Proc Natl Acad Sci U S A 112, 11072–11077, doi:10.1073/pnas.1504242112 (2015).

83 Northoff, G. & Bermpohl, F. Cortical midline structures and the self. Trends Cogn Sci 8, 102–107, doi:10.1016/j.tics.2004.01.004 (2004).

84 Summerfield, J. J., Hassabis, D. & Maguire, E. A. Cortical midline involvement in autobiographical memory. Neuroimage 44, 1188–1200, doi:10.1016/j.neuroimage.2008.09.033 (2009).

85 Svoboda, E., McKinnon, M. C. & Levine, B. The functional neuroanatomy of autobiographical memory: a meta-analysis. Neuropsychologia 44, 2189–2208, doi:10.1016/j.neuropsychologia.2006.05.023 (2006).

86 Wilson, B. A. et al. Egocentric disorientation following bilateral parietal lobe damage. Cortex 41, 547–554, doi:10.1016/s0010-9452(08)70194-1 (2005).

87 Ciaramelli, E., Rosenbaum, R. S., Solcz, S., Levine, B. & Moscovitch, M. Mental space travel: damage to posterior parietal cortex prevents egocentric navigation and reexperiencing of remote spatial memories. J Exp Psychol Learn Mem Cogn 36, 619–634, doi:10.1037/a0019181 (2010).

88 Whitlock, J. R., Pfuhl, G., Dagslott, N., Moser, M. B. & Moser, E. I. Functional split between parietal and entorhinal cortices in the rat. Neuron 73, 789–802, doi:10.1016/j.neuron.2011.12.028 (2012).

89 Wilber, A. A., Clark, B. J., Forster, T. C., Tatsuno, M. & McNaughton, B. L. Interaction of egocentric and world-centered reference frames in the rat posterior parietal cortex. J Neurosci 34, 5431–5446, doi:10.1523/JNEUROSCI.0511-14.2014 (2014).

90 Sargolini, F. et al. Conjunctive representation of position, direction, and velocity in entorhinal cortex. Science 312, 758–762, doi:10.1126/science.1125572 (2006).

91 Slater, M., Spanlang, B., Sanchez-Vives, M. V. & Blanke, O. First person experience of body transfer in virtual reality. PLoS One 5, e10564, doi:10.1371/journal.pone.0010564 (2010).

92 Sanchez-Vives, M. V., Spanlang, B., Frisoli, A., Bergamasco, M. & Slater, M. Virtual hand illusion induced by visuomotor correlations. PLoS One 5, e10381, doi:10.1371/journal.pone.0010381 (2010).

93 Weech, S., Kenny, S. & Barnett-Cowan, M. Presence and Cybersickness in Virtual Reality Are Negatively Related: A Review. Front Psychol 10, 158, doi:10.3389/fpsyg.2019.00158 (2019).

94 Fischl, B. & Dale, A. M. Measuring the thickness of the human cerebral cortex from magnetic resonance images. Proc Natl Acad Sci U S A 97, 11050–11055, doi:10.1073/pnas.200033797 (2000).

95 Desikan, R. S. et al. An automated labeling system for subdividing the human cerebral cortex on MRI scans into gyral based regions of interest. Neuroimage 31, 968–980, doi:10.1016/j.neuroimage.2006.01.021 (2006).

96 Stangl, M., Shine, J. & Wolbers, T. The GridCAT: A Toolbox for Automated Analysis of Human Grid Cell Codes in fMRI. Front Neuroinform 11, 47, doi:10.3389/fninf.2017.00047 (2017).

97 Eickhoff, S. B., Heim, S., Zilles, K. & Amunts, K. Testing anatomically specified hypotheses in functional imaging using cytoarchitectonic maps. Neuroimage 32, 570–582, doi:10.1016/j.neuroimage.2006.04.204 (2006).

