## Supplementary Figure for "Sense of self impacts spatial navigation and hexadirectional coding in human entorhinal cortex"

### Supplementary Information

#### Supplementary Text

##### Supplementary Results

###### Detailed analysis on the subparameters of GCLR

We further assessed the subparameters of GCLR (i.e. voxel-wise amplitude, temporal/spatial stability of the grid orientation) to better understand the mechanisms underlying the decreased GCLR we observed in the Body condition. Thus, to better probe the magnitude of the BOLD activity contrast that might possibly be occluded by unstable grid orientations, we averaged voxel-wise amplitudes of the estimated six-fold sinusoidal curve in the EC region of interest (ROI) regardless of the grid-orientation value the voxel has. The voxel-wise amplitudes were calculated to estimate mean grid-orientation during the conventional GCLR analysis, and have been regarded to reflect grid cell-like activity at the voxel level (i.e. the values are used as weighting factors while calculating the mean grid orientation in the ROI)[1,2]. Comparing the mean voxel-wise amplitudes across conditions, we replicated our finding that the values were significantly lower in the Body vs. No-body condition (Z = 2.68, r = 0.54, p = 7.37e-03, n = 25; Supplementary Figure 4a). We also found a significant correlation between the mean voxel-wise amplitudes of the hexadirectional modulation and the condition-wise GCLRs (linear mixed-effects regression; NumDF = 1, DenDF = 142, F = 4.58, p = 0.034; Supplementary Figure 4d). Notably, the values were even more strongly correlated to the absolute values of the GCLR (NumDF = 1, DenDF = 135.9, F = 20.69, p = 2.22e-03; Supplementary Figure 4g), consistent with the theoretical expectation that a sign of GCLR can be inverted to negative by wrongly estimated grid cell orientation, although the specific BOLD modulation itself is prominent. The spatial stability of grid orientations, defined as the homogeneity of voxel-wise grid orientations within the EC-ROI, was quantified by Rayleigh's Z (see Methods)[3,4]. In addition, to quantify temporal stabilities of the grid orientations during a session, standard deviations of grid orientations estimated from different portions of the session were calculated (indexing instability rather than stability; see Method). These analyses revealed that grid orientations in the Body condition were both spatially and temporally less stable, although these were not found to significantly differ between both conditions (Spatial stability: Z = 1.25, r = 0.25, p = 0.210; Temporal instability: Z = 1.47, r = 0.29, p = 0.141; Supplementary Figure 4b-c). We note, further, that both spatial and temporal stability, significantly influenced the condition-wise GCLRs (NumDF = 1, DenDF = 142, Spatial stability: F = 9.27, p = 2.78e-03;

Temporal instability: F = 23.81, p = 2.82e-06; Supplementary Figure 4e-f).

#### Supplementary Discussion

##### Participant's position

Humans rarely navigate in a supine position and it could be argued that the decreased GCLR we observed is related to such mechanisms, potentially enhanced by showing a supine avatar during navigation. However, several arguments speak against this hypothesis. First, the participants were in a supine position in both conditions. Second, this was the same position as tested in all previous human GCLR work using fMRI. Third, as previous spatial navigation work did not quantify experienced self-location and self-identification with the avatar, the experienced position of participants for previous GCLR data is, therefore, to the best of our knowledge, not known. We believe that although humans mostly navigate in an upright position, especially the sitting position (e.g. car, wheelchair) and also the supine position (e.g. luge) are suitable positions to test spatial navigation. More work is necessary for this important issue. The present data suggest that for any spatial navigation paradigm (upright, sitting, supine) the control of the different bodily reference frames (i.e. somatosensory, visual), as well as the quantification of the subjective bodily reference frame (i.e. self-identification, self-location), should be controlled.

##### Drift in self-location

It could be also interesting to assess whether the drift in experienced self-location that we observed in the Body condition (Fig. 3) affected the grid cell pattern (e.g. shift of hexagonal grid). However, our GCLR analyses were based on the seminal work by Doeller and colleagues (2010), who based GCLR on the analysis of the heading direction-dependent hexadirectional BOLD modulation, which is calculated with heading direction information that is independent of the individual's location in the arena. Hence, as the small drift in self-location (~1vm) that we report was along the same heading direction, there can be no change in GCLR (i.e. a heading-direction-dependent BOLD modulation). Further dedicated experiments would be required, inducing drifts in self-location, that possibly generate or affect detectable GCLR metrics.

**Other parameters that might influence on GCLR**

Rodent single neuron recordings and human GCLR results have demonstrated that grid cell activity depends on the navigation speed of the subject[1,5,6]. However, speed differences cannot account for the present GCLR reduction, because navigation velocity was fixed not to differ between both conditions. Moreover, considering that the navigated distance was shorter in the Body vs. No-body condition (with the same navigation time) a speed-related effect should have rather led to higher, not lower, GCLR in the Body condition. The average moving speeds of each trial (navigated distance/time), indeed, did not differ between conditions ($p = 0.46$).

In addition, although we report a larger distance from the border in the Body condition, further analysis revealed that the central navigational preference (proposed by Kunz and colleagues, 2015) did not differ in the two conditions of the present experiment ($p = 0.84$). The undershot (drift in self-location) we observed only happened at the last moment of each navigation and was quite subtle (~1 vm on average) compared to the much larger diameter of the arena (110 vm). The decreased GCLR can also not be explained by the difference in target locations as all objects and target locations were randomly shuffled across conditions. Additional analysis also showed that head motion artifacts did not differ between conditions (Body: 0.240±0.012 mm, No-body: 0.247±0.014 mm; $p = 0.08$) and were taken into account by the nuisance regressors with the motion parameters.

As participants were only trained in the No-body condition, it could be argued that previous exposure to the No-body, but not the Body condition, may account for the observed GCLR differences between conditions (i.e. re-exposure effect). This, however, does not seem very likely, because when reanalyzing our GCLR results across conditions, but by excluding the 1st block for each condition, we confirmed our original results and found that only the GCLR in the No-body condition was significant ($p = 6.23e-3$), whereas this was not the case in the Body condition ($p = 0.28$).

Finally, it could be argued that the reduced GCLR during the Body condition is related to distraction or visual occluding of the VR scene by the avatar. However, as shown by the exemplary task scenes (Fig. 1; see also supplementary video1), there was only minimal spatial information that participants could acquire from the lower part of the scene (occluded by the avatar) as all landmarks were placed in a distal position and not in the task arena, where they were navigating. It is, therefore, unlikely that the present effects were caused by scene occlusion due to the avatar. Moreover, distraction or occlusion (due to the avatar) should lead to decreases in spatial navigation performance and we observed the opposite effect: spatial navigation performance was better in the Body condition.

#### Hippocampal BOLD activity and compensatory mechanisms

Kunz et al. (2015) described that activation in the hippocampus might serve as a compensatory mechanism in cases where GCLR was found to be attenuated. It could therefore be argued that the present changes in GCLR across conditions may reflect similar compensatory hippocampal activity. However, hippocampal BOLD activity did not differ in the Body versus No-body condition in the present study ($p = 0.94$).

### Supplementary FIGURES

#### Supplementary Figure 1

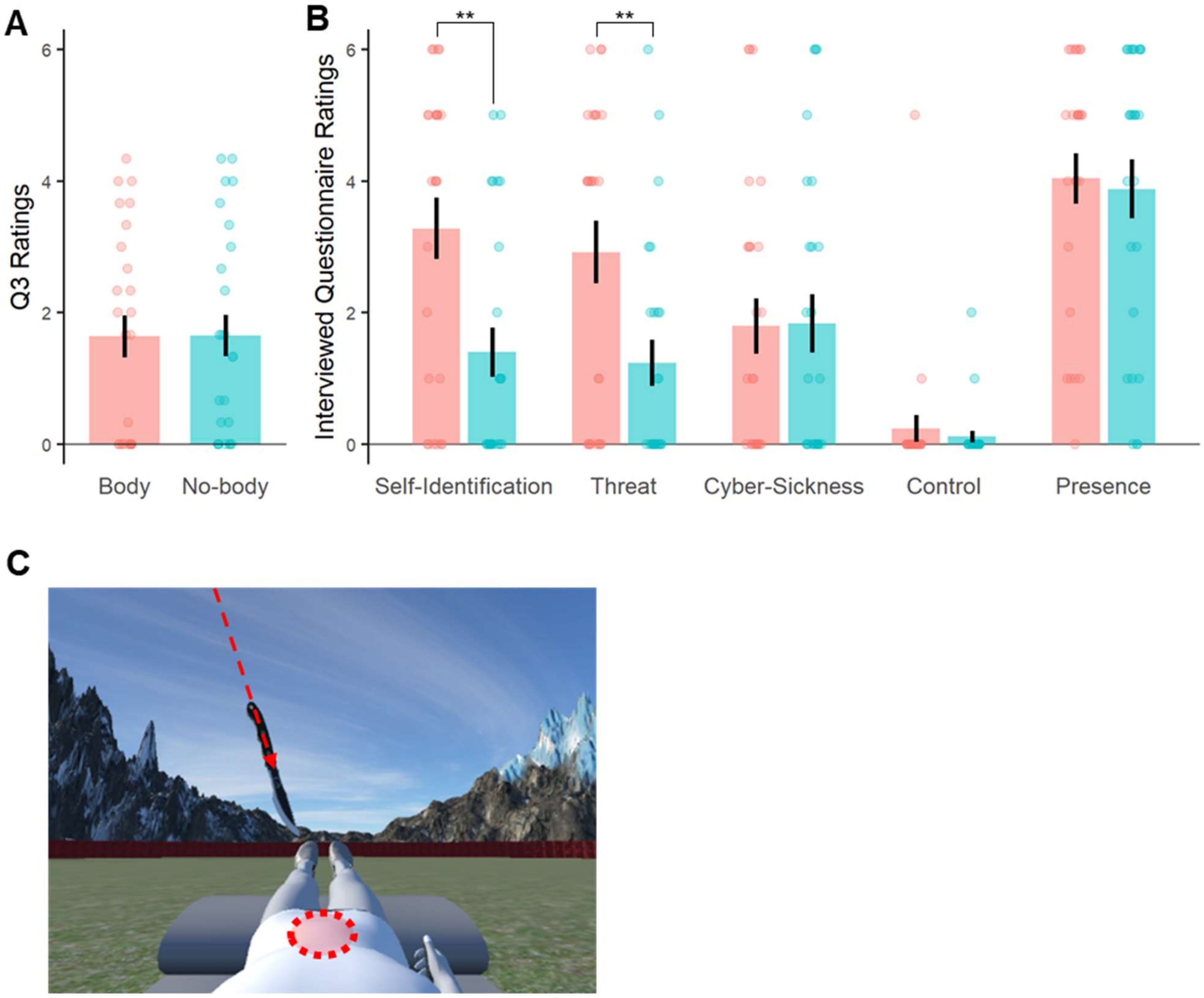


**Fig. S1, related to Fig. 2 and the questionnaire section of the method. Question3 (Q3) ratings and questionnaire ratings from the post-experiment interviews. (A)** Q3 ratings capturing cyber-sickness of participants did not differ between the experimental conditions ($p = 0.57$) **(B)** Verbally answered questionnaire ratings from the post-experiment interviews. The results showed the difference in ‘self-identification’ and ‘Threat’ ratings between the experimental conditions, which is in accordance with the questionnaire results answered during the task. A question for ‘Presence (i.e. spatial immersion into the virtual space’) was included in the post-experiment interview and did not statistically differ between the two experimental conditions. **(C)** The virtual threat (i.e. knife) was directed to the position as indicated by the red oval (red oval was not shown) in both conditions. ** : $p < 0.01$

#### Supplementary Figure 2

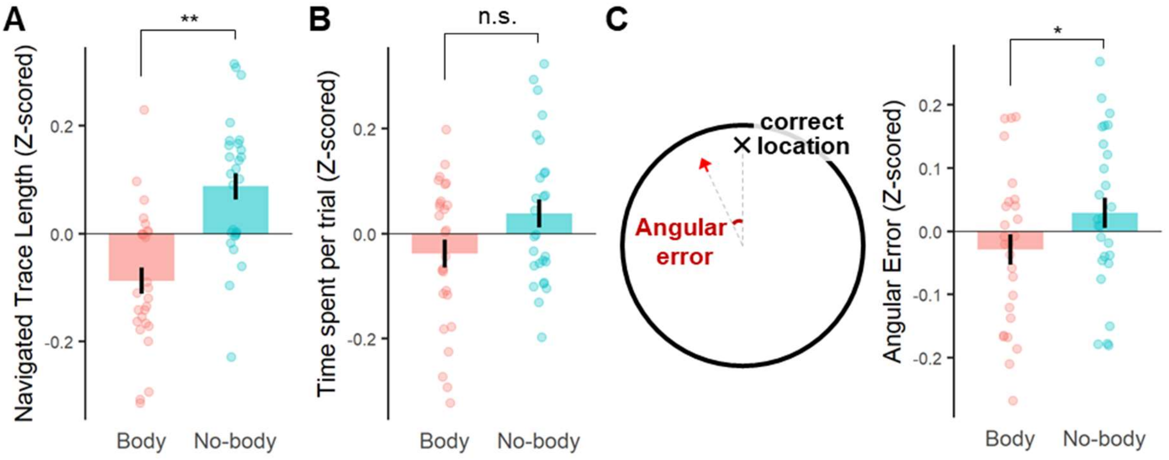


**Fig. S2, related to Fig. 2,3. Spatial navigation efficiency and spatial memory precision independent of the self-drift effect were better in the Body condition (A)** The participants navigated through significantly shorter traces with the embodied avatar **(B)** while they spent similar time during the trial. **(C)** Angular errors calculated regardless of the distance from the border were compared between the two experimental conditions. The results consistently showed that spatial memory precision was better (indicated by the lower angular errors) in the Body condition. n.s. : $p >= 0.05$, * : $0.01 <= p < 0.05$, ** : $p < 0.01$

**Supplementary Figure 3**

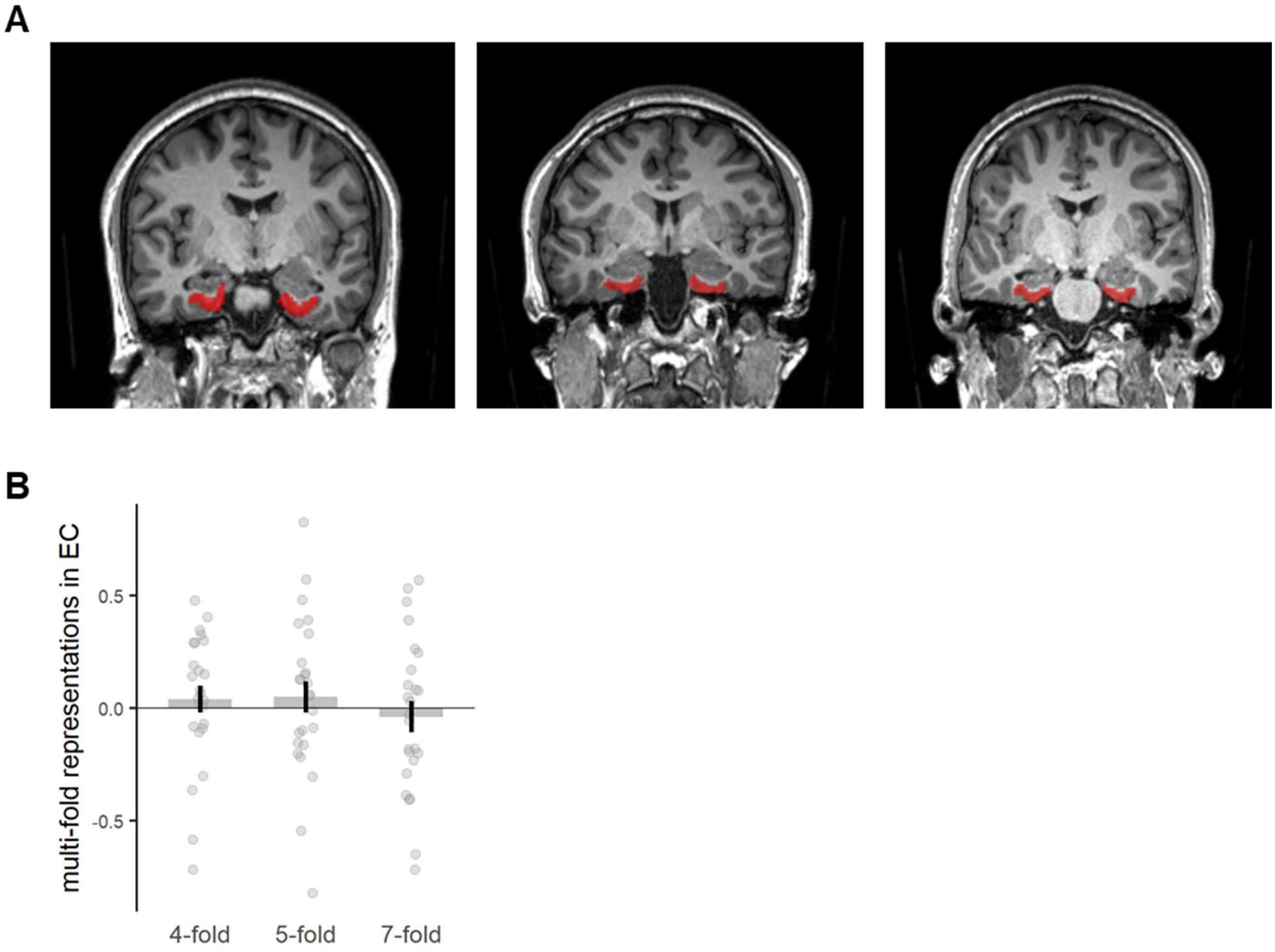


**Fig. S3, related to Fig. 4. (A)** EC ROIs from 3 Exemplary participants, **(B)** Control multi-fold representations were not significantly greater than zero.

#### Supplementary Figure 4

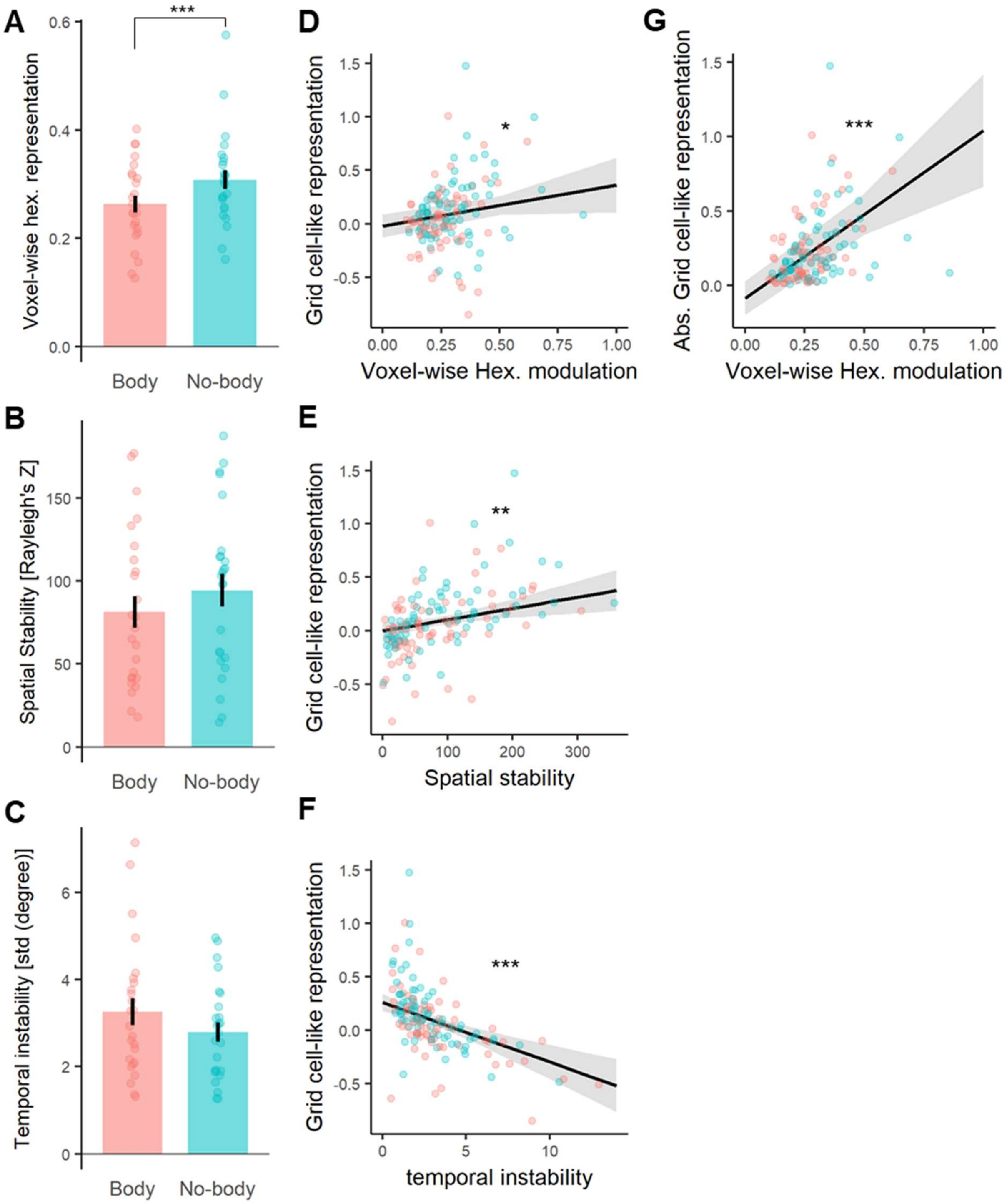


**Fig. S4, related to Fig. 5. Parameters relevant to estimating Grid cell-like representation (GCLR) were consistently better in the Body condition. (A)** Voxel-wise hexadirectional modulation in the entorhinal cortex(EC) was significantly attenuated in the Body condition. **(B)** Spatial Stability (i.e. homogeneity of voxel-wise grid orientations in EC) was also lower in the Body condition, but the difference did not reach significance. **(C)** Grid orientations were less stable in time, during the Body condition. However, the difference was insignificant. **(D,E,F)** Both the voxel-wise amplitude of the hexadirectional modulation and the spatial stability were significantly and positively correlated with the estimated GCLR, while temporal instability was negatively correlated. Contributions of the three parameters(voxel-wise amplitude, spatial stability, and temporal instability) to the estimated GCLR was assessed simultaneously with a multiple mixed-effect model. **(G)** As theoretically expected, the voxel-wise amplitude of the six-fold modulation was more strongly correlated with the absolute values of GCLRs, which were less affected by wrongly calculated grid orientations. Of note, figure A-C are plotted with subject-wise mean values, while figure D-G are plotted with session-wise mean values; no data points were excluded. This was based on the statistical analysis that we applied on the data depicted respectively in each figure: 1) a Wilcoxon's signed-rank test to assess the condition-wise difference and 2) a mixed effect model to assess the correlation between metrics with every round-wise value. * : $0.01 <= p < 0.05$, ** : $0.001 <= p < 0.01$, *** : $p < 0.001$.

#### Supplementary Figure 5

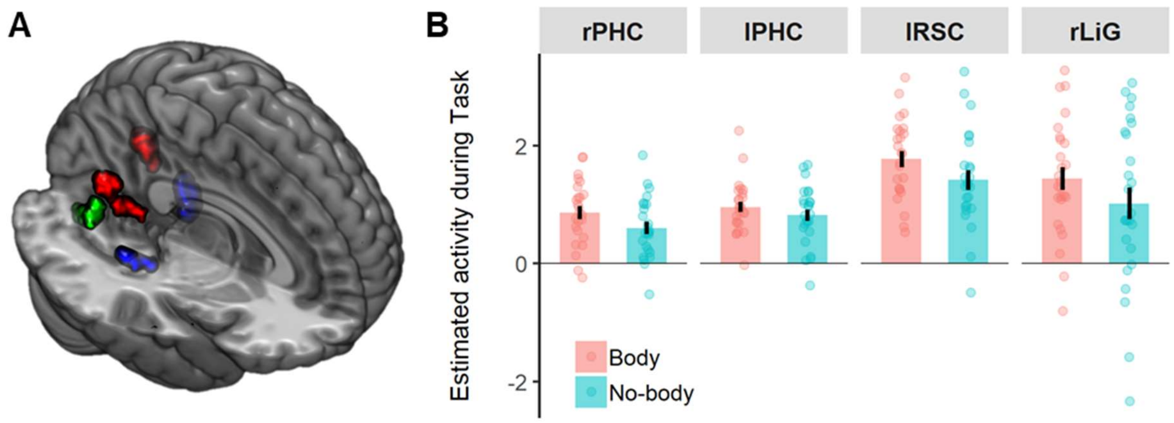


**Fig. S5, related to Fig. 6. Task-related brain regions defined by the functional localizer.**

**(A)** The bilateral parahippocampal gyrus (PHC: Blue), bilateral retrosplenial cortex (RSC: Red; see Fig. 6 for the right RSC), and right lingual gyrus (rLiG: Green) were activated during the spatial navigation task procedure. **(B)** In the four other brain regions revealed by the analysis (except for the right RSC), we could not find a significant difference between the conditions.

#### Supplementary Tables

| Regressor | Parametric modulator | Duration per trial |
|---|---|---|
| Cue | | 2.5 s |
| Retrieval | | variable |
| Retrieval | Distance error | variable |
| Self-Estimation | | variable |
| Feedback | | 2.0 s |
| Collection | | variable |
| Threat | | 2.5 s |

**Supplementary Table 1.** List of regressors for whole-brain GLM analysis

| Region | MNI coordinates (mm) | Cluster Size (voxels) | Peak t-value | Peak p-value |
|---|---|---|---|---|
| **Retrosplenial cortex / Precuneus** | | | | |
| Right | 20, -56, 22 | 121 | 10.67 | <0.001 |
| Left | -16, -60, 22 | 114 | 11.20 | <0.001 |
| **Parahippocampal gyrus** | | | | |
| Right | 24, -40, -10 | 58 | 13.21 | <0.001 |
| Left | -22, -42, -8 | 125 | 12.34 | <0.001 |
| **Lingual gyrus** | | | | |
| Right | 8, -70, -2 | 90 | 9.49 | <0.001 |

$p < 0.05$, whole-brain, voxel-wise FWE correction

**Supplementary Table 2.** Task-related brain regions established by the functional localizer

#### Legend for Supplementary Movie S1

**NavigationTask_Moon_et_al.mp4**

: The movie shows a participant's physical body (focusing on the right hand) in the scanner and the corresponding task scene while they were performing the virtual navigation task in the Body condition.

#### SI References


1 Doeller, C. F., Barry, C. & Burgess, N. Evidence for grid cells in a human memory network. *Nature* **463**, 657-661, doi:10.1038/nature08704 (2010).

2 Stangl, M., Shine, J. & Wolbers, T. The GridCAT: A Toolbox for Automated Analysis of Human Grid Cell Codes in fMRI. *Front Neuroinform* **11**, 47, doi:10.3389/fninf.2017.00047 (2017).

3 Kunz, L. *et al.* Reduced grid-cell-like representations in adults at genetic risk for Alzheimer's disease. *Science* **350**, 430-433, doi:10.1126/science.aac8128 (2015).

4 Stangl, M. *et al.* Compromised Grid-Cell-like Representations in Old Age as a Key Mechanism to Explain Age-Related Navigational Deficits. *Curr Biol* **28**, 1108-1115 e1106, doi:10.1016/j.cub.2018.02.038 (2018).

5 Sargolini, F. *et al.* Conjunctive representation of position, direction, and velocity in entorhinal cortex. *Science* **312**, 758-762, doi:10.1126/science.1125572 (2006).

6 Kropff, E., Carmichael, J. E., Moser, M. B. & Moser, E. I. Speed cells in the medial entorhinal cortex. *Nature* **523**, 419-424, doi:10.1038/nature14622 (2015).
